# In vivo ephaptic coupling allows memory network formation

**DOI:** 10.1101/2023.02.28.530474

**Authors:** Dimitris A. Pinotsis, Earl K. Miller

## Abstract

It is increasingly clear that memories are distributed across multiple brain areas. Such “engram complexes” are important features of memory formation and consolidation. Here, we test the hypothesis that engram complexes are formed in part by bioelectric fields that sculpt and guide the neural activity and tie together the areas that participate in engram complexes. Like the conductor of an orchestra, the fields influence each musician or neuron and orchestrate the output, the symphony. Our results use the theory of synergetics, machine learning and data from a spatial delayed saccade task and provide evidence for in vivo ephaptic coupling in memory representations.

## Introduction

In recent decades there has been a paradigm shift in neuroscience. In the past, we focused on properties of individual neurons(James, 1890; Queenan et al., 2017). There is now a growing realization that information storage and processing depends on spatially distributed, dynamic groupings of neurons (Buschman et al., 2011; Fujisawa et al., 2008; Yuste, 2015), known as neural ensembles (Buschman et al., 2012; Pfau et al., 2013; Pinotsis et al., 2017a; Pinotsis and Miller, 2017; Tayler et al., 2013) or engram cells (Josselyn et al., 2015; Thompson, 1976). Techniques like protein induction (Gordon et al., 1980), immediate early gene (IEG) expression (Guzowski et al., 2005) and optogenetics (Fenno et al., 2011) allow for identification of ensemble neurons participating in memory storage and recall (Ryan et al., 2015; Tonegawa et al., 2015b). Further, recent experiments have found simultaneous neural ensembles maintaining the same memory in many brain areas, something known as engram complex (Poo et al., 2016; Roy et al., 2019). In (Roy et al., 2019) a total of 247 brain areas were mapped using the protein cFos and IEG. Among them, 117 areas were found to be significantly reactivated when a fear memory was recalled. Thus, memory was not stored in a single brain area but was dispersed in multiple areas and neural ensembles. Earlier theories like memory consolidation(Squire and Alvarez, 1995) and multiple traces (Nadel and Moscovitch, 1997) have also found that memories are stored in multiple areas forming engram complexes. These are connected via engram pathways formed by mono- or poly-synaptic connections (Tonegawa et al., 2015a).

The challenge, then, is in understanding how the brain forms engram complexes. Each brain area is connected to many others. Anatomical connectivity alone cannot be the whole story. Hypotheses that could explain this include that engram complexes are dynamically formed by emergent properties of neurons like synchronized rhythms (Harris et al., 2003; Lundqvist et al., 2018; Miller et al., 2018, 2014), possibly resulting from internal coordination of spike timing (Koch, 2004; Singer, 1999), that allow neuronal communication (Fries, 2015; Lakatos et al., 2019; Reinhart and Nguyen, 2019), feature integration and perceptual segmentation (Engel and Singer, 2001; Moore and Obhi, 2012). Here, we report tests of the hypothesis that the electric fields generated by neurons play a crucial role. We suggest that ephaptic coupling (Anastassiou et al., 2011; Ruffini et al., 2020) ties together the areas that participate in engram complexes. In other words, we test the hypothesis that memory networks include electric fields that carry information back to individual neurons.

Direct evidence of ephaptic coupling of spiking has been found in brain slices (Anastassiou and Koch, 2015; Chiang et al., 2019; Jefferys et al., 2012). *In vitro* ephaptic coupling has been found in LFPs. Application of external electric fields resulted in membrane potentials oscillating at the same frequency as the drive(Anastassiou et al., 2011). Support for its role in forming engram complexes comes from studies showing that neurons participating in an engram complex showed similar functional connectivity during optogenetic activation and memory recall(Roy et al., 2019),(Kitamura et al., 2017). We found that the electric fields in the primate prefrontal cortex carried information about the contents of working memory (Pinotsis and Miller, 2022a). Using data from a delayed saccade task (Jia et al., 2017; Pinotsis et al., 2017a), we built two models: one for neural activity (Pinotsis et al., 2017a; Pinotsis and Miller, 2017) and another for the emergent electric field. This revealed electric field patterns that varied with contents of working memory. Further, we found that the electric fields were robust and stable while neural activity underlying memory showed representational drift. This latter observation suggested the hypothesis that electric fields could act as “guard rails” that help stabilize and funnel the high dimensional variable neural activity along stable lower-dimensional routes.

Here we test the hypothesis that electric fields sculpt and guide the neural activity forming engram complexes. We used a theory of complex systems known as synergetics (Haken, 2012, 1987). We also extended the single area analysis of (Pinotsis and Miller, 2022a) and focused on data from two areas known to form an engram complex, Frontal Eye Fields (FEF) and Supplementary Eye Fields (SEF). Synergetics describes how complex systems (e.g. molecules, fluids, brain etc.) self – organize. In the case of human behavior, synergetics describe how the collective dynamics of muscles and body parts (e.g. fingers) give rise to behavior like rhythmic hand movement (Haken et al., 1985). We applied synergetics to understand the emergence of memory representations. We performed mathematical, i.e. pen and paper, computations and showed that the theory predicts that electric fields guide ensemble activity. If ephaptic coupling occurs in a brain area and this exchanges memory information with other brain areas, then ephaptic coupling will occur in those areas too. We then confirmed our results using Bayesian Model Comparison (Friston and Penny, 2011; Kass and Raftery, 1995), Granger Causality (Barnett and Seth, 2014) and Representation Similarity Analysis (Kriegeskorte et al., 2008). This suggested that the electric field enslaves neurons, not the other way around. Applying the slaving principle (Haken, 2012), we found that the electric field controls neural activity and oscillations through ephaptic coupling (Anastassiou and Koch, 2015; Fröhlich and McCormick, 2010) and that this was the case across all recording sites that participated in the engram complex.

## Methods

### Mathematical Notation

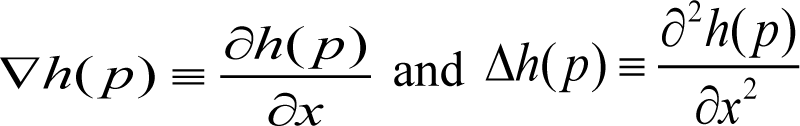 (first and second derivatives evaluated at point *p*), (*j*) ∂ *^j^h*(*x*,*t*) for an arbitrary function *h*. 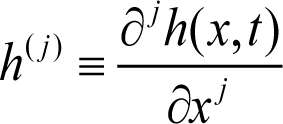 denotes the spatial derivative of order *j*. The subscript “0” denotes boundary values, e.g. Δ*V*_0_^*e*^ is the value of the second derivative of the extracellular potential *V ^e^* on the exterior of the membrane. A random process *Ṽ^m^* from which the transmembrane potential *V ^m^* is sampled is denoted by tilde with samples *Ṽ^ml^*, indexed by *l*. Hat denotes the Fourier Transform (FT) of a function *h*, i.e. 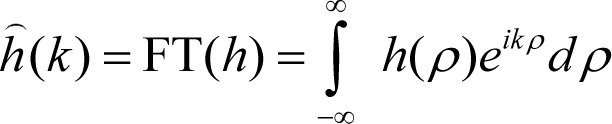.

### Task and Experimental Setup

We reanalyzed data from (Jia et al., 2017). The same data were used in our earlier papers (Pinotsis et al., 2017b; Pinotsis and Miller, 2022b). Two adult male Macaca monkeys were trained to perform an oculomotor spatial delayed response task. This task required the monkeys to maintain the memory of a randomly chosen visual target (at angles of 0, 60, 120, 180, 240 and 300 degrees, 12.5-degree eccentricity) over a brief (750 ms) delay period and then saccade to the remembered location. If a saccade was made to the cued angle, the target was presented with a green highlight and a water reward was delivered. If not, the target was presented with a red highlight and reward was withheld. 32-electrode chronic arrays were implanted unilaterally in FEF and SEF in each monkey. Each array consisted of a 2 x 2 mm square grid, where the spacing between electrodes was 400 um. The implant channels were determined prior to surgery using structural magnetic resonance imaging and anatomical atlases. From each electrode, we acquired local field potentials (LFPs; extracted with a fourth order Butterworth low-pass filter with a cut-off frequency of 500Hz, and recorded at 1 kHz) using a multichannel data acquisition system (Cerebus, Blackrock Microsystems). We analyzed LFPs during the delay period when monkeys held the cued angles in memory.

### A neural field model of ephaptic coupling

This section summarizes a theoretical model for the description of neural ensemble activity developed earlier (Pinotsis et al., 2017; Pinotsis and Miller, 2022, 2017). We modelled the activity of neural ensembles. These are groups of neurons that maintain working memory representations. Some results about their activity summarized below involve lengthy derivations not repeated here. The interested reader might consult earlier papers that are referenced and the *Supplementary Material*.

In earlier work, we used neural field theory (Coombes, 2005; Deco et al., 2008; Jirsa and Haken, 1996; Robinson et al., 2016), (cf. Equation (4) in (Pinotsis et al., 2017)) to describe the evolution of the transmembrane potential or depolarization, *V^m^*, in neural ensembles. Currents flow along the neurons’ axons and dendrites. Chemical energy is converted to electrical. Action and synaptic potentials are summed up to produce an emerging *electric potential* (EP) *V^e^* in extracellular space. The difference of intracellular *V ^i^* and extracellular *V^e^* potentials on either side of the membrane, *V^m^* = *V^e^*_0_ − *V^i^*_0_ is the transmembrane potential (recall that the subscript “0” denotes boundary values). The time evolution of the transmembrane potential *V^m^* can be described by a neural field model (Atay and Hutt, 2004; Bojak et al., 2013; Pinotsis et al., 2012). Figure 1A includes a schematic of a chronic array implanted in a cortical area (for simplicity, 10 electrodes shown as dots in the blue square). Each electrode is thought to be sampling from a neural population in its proximity and we assumed that the ensemble occupies a patch (cortical manifold) denoted by *Δ*. Activity is sampled at the locations of the electrodes. It is thought to be generated by a neural population in the vicinity of the electrode. To construct the neural field model, we numbered the electrodes in a monotonic fashion (cf. the numbers in Figure 1A). For mathematical convenience, we also assumed that *Δ* can be replaced by a line, i.e. electrodes are all next to each other (cf. red line at the bottom of Figure 1A). This assumption was tested in (Pinotsis et al., 2017) and (Pinotsis and Miller, 2022a). There, we found that the model explained more than 40% of the data variance. A second test of this assumption is discussed after Equation (3) below. The coloured curves connecting electrodes (dots on the red line at the bottom of Figure 1A) are schematics of Gaussian functions that describe connectivity between electrodes and populations, see (Pinotsis et al., 2017) for details.

**Figure 1.**
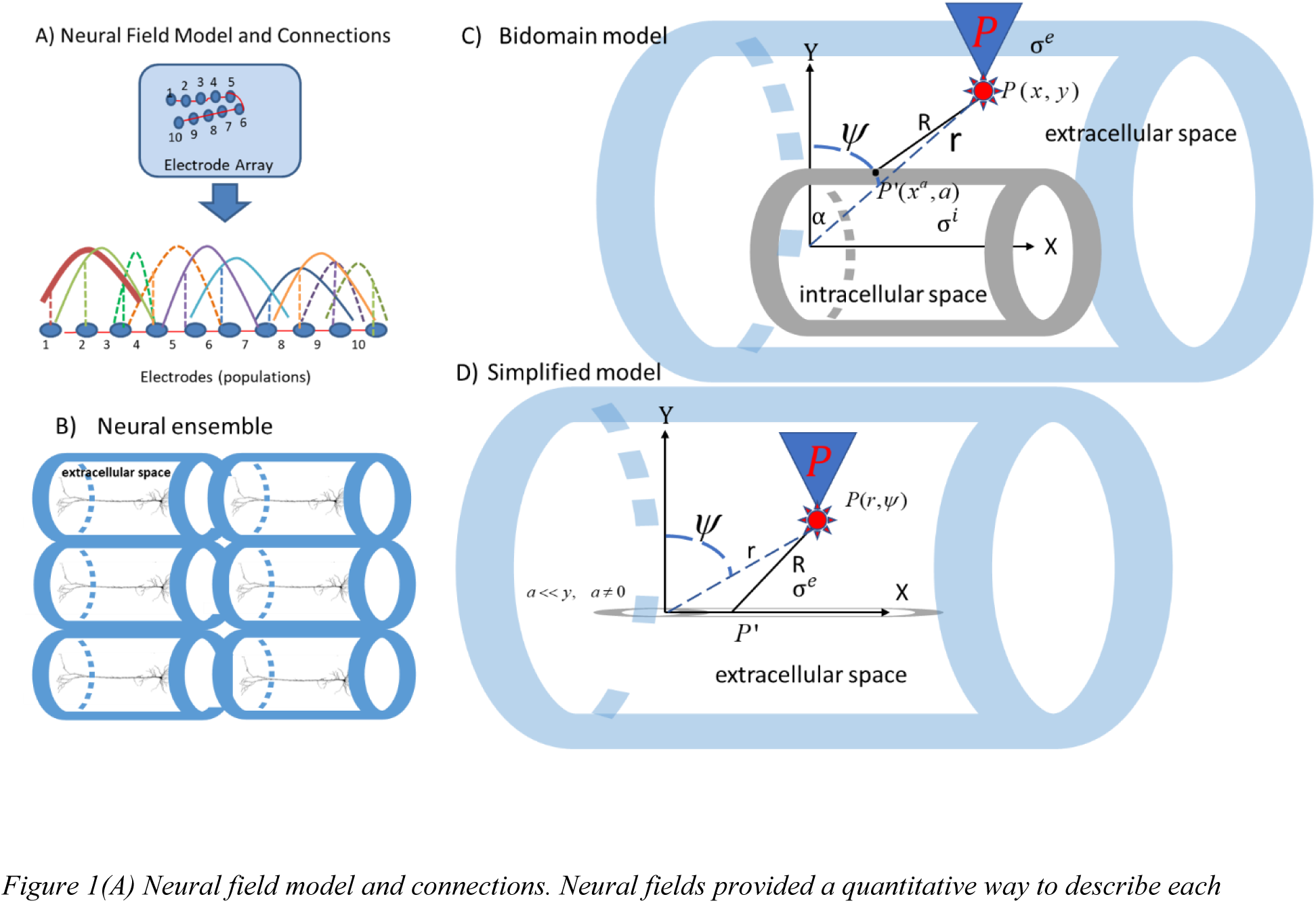
(A) Neural field model and connections. Neural fields provided a quantitative way to describe each ensemble’s patterns of activity across simultaneously recorded sites. The same model can describe different ensembles. Each electrode occupies a position on a cortical manifold (line) Δ parameterized by the variable x and is connected to all other electrodes with connections whose strength follows a Gaussian profile (coloured solid and dashed lines), see (Pinotsis et al., 2017) for more details. (B) Extracellular space around each neuron within the ensemble (blue cylindrical fibers). (C) Bidomain model for the electric field generated by a cylindrical fiber in a conductor. The extracellular and intracellular space are depicted by blue and grey cylindrical fibers (see Methods for the meaning of various symbols). (D)Simplified bidomain model where the measurement point is located at a vertical distance much larger than the radius of intracellular space.

Our neural field model describes transient fluctuations around baseline, similar to spontaneous activity in large scale resting state networks (Deco et al., 2010; Drysdale et al., 2017; D.A. Pinotsis et al., 2013). It predicts average firing rate or depolarization, similar to activation functions in deep neural networks (LeCun et al., 2015; Pinotsis et al., 2019).

Mathematically, the neural field model suggests that the time evolution of depolarization *V^m^* is given by the following equation (see also Equation (4) in (Pinotsis et al., 2017)):

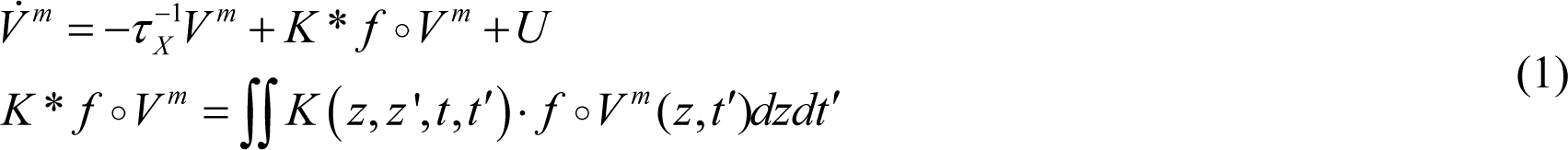

Equation (1) suggests that *V^m^* changes as a result of three terms: a simple decay, recurrent inputs from other parts of the ensemble and some exogenous, stochastic input *U.* We called this neural field “*deep*” to distinguish this model (with learned connectivity parameters) from common neural field models where connectivity weights are chosen *ad hoc*. The integral appearing in Equation (1) is defined over the cortical patch, i.e. *z* ϵ Δ and *t* > 0. It describes how the diffusion of local recurrent input changes *V^m^*. Here, *z* parameterizes the location on a cortical patch occupied by the ensemble, *X* is an index denoting excitatory or inhibitory populations, *K* is the connectivity or weight matrix that describes how the signal is amplified or attenuated when it propagates between electrodes (cf. coloured curves in Figure 1A), *U* is endogenous neural input and 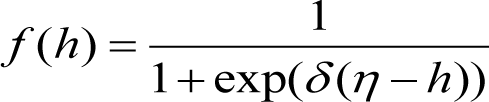 is called transfer function. Also, *τ _X_* is the time constant of postsynaptic filtering, *δ* is synaptic gain and *η* is the postsynaptic potential at which the half of the maximum firing rate is achieved, see e.g. (Pinotsis et al., 2012) for more details.

In (Pinotsis et al., 2017), we assumed that the transmembrane potential *V^m^_X_* is sampled from a random process *Ṽ^m^* with samples 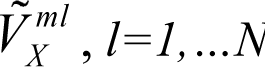. We then considered a new variable 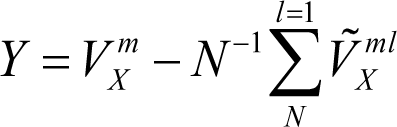 and showed that Equations (1) can be reformulated as a Gaussian Linear Model (GLM)

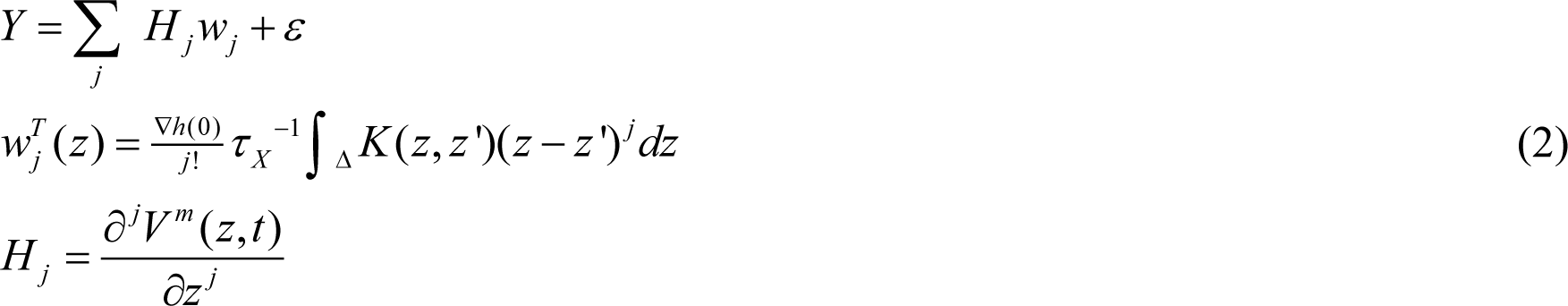

where *ε* (*m*, *s*_*s*_^2^ *I*) and *s_s_* is the inverse precision. Note that *m* is the sample mean 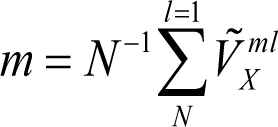. For a detailed derivation of Equation (2) from Equation (1) and its relation to similar models like Wilson and Cowan (Wilson and Cowan, 1973), see (Pinotsis and Miller, 2022a) and *Supplementary Material*. In Equation (2), the functions *w* are called the connectivity components and *H* are the principal axes. The connectivity components *w* (second line in Equation (2)) provide the connectivity matrix *K* (cf. Equation (1) and Figure 1A). They describe how signal recorded from a certain electrode contributes to LFP data (across all trials). They are of dimensionality number of electrodes *by* number of trials. The principal axes (last line in Equation (2)) are matrices of dimensionality number of time samples *by* number of trials. They describe the average instantaneous contribution to the LFP data across all electrodes. Please see (Pinotsis et al., 2017; Pinotsis and Miller, 2022) as well as the *Supplementary Material* for more details about and the connectivity components and the principal axes.

To find the connectivity components *w,* we used a Restricted Maximum-Likelihood (ReML) algorithm (Harville, 1977a). This optimizes a cost function known as the Free Energy, *F*,

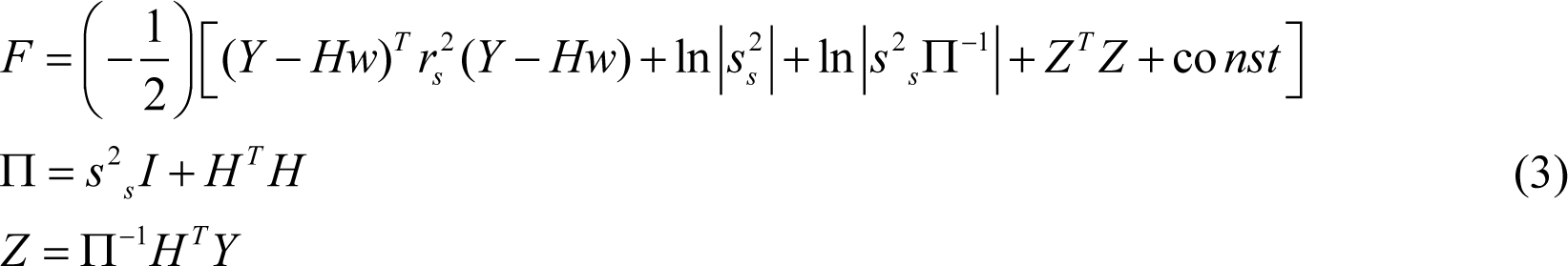

The connectivity *w* was obtained by training the neural field model given by Equation (1) using the cost function given by Equation (3) maximises the mutual information between the remembered cue and the ensemble activity. This can be thought to describe synaptic efficacy in a neural ensemble that represents a certain stimulus. In (Pinotsis and Miller, 2022) we obtained the connectivity matrices and compared them to the connectivity obtained using two independent methods: *k*-means clustering (Humphries, 2011) and high dimensional SVD (Carroll and Chang, 1970; Williams et al., 2018). This served as a validation of the neural field model given by Equation (1). It also provided a second test of the earlier assumption where we replaced the cortical patch Δ by a line (red line at the bottom of Figure 1A). We found that the connectivity obtained after training the neural field model with the cost function (3) is the same as the connectivity found using pairwise correlations (Humphries, 2011) and SVD (Williams et al., 2018).

To sum up, in previous work we showed that neural fields given by Equations (1) can be rewritten like a Gaussian Linear Model given by Equation (2). We also showed how neural fields can be trained using the Free Energy given by Equation (3) to obtain the connectivity *K*. In (Pinotsis and Miller, 2022) we also showed that Wilson and Cowan network models (Wilson and Cowan, 1973) can also be written in the form of a GLM and trained using the cost function (3). Here, we will use neural field models given by Equation (1).

Below, we will consider an extension of the model (1) that will include ephaptic coupling (interactions between emerging electric fields produced by neural ensembles and the underlying neural activity). Later, we will fit this extended as well as the original neural field model to LFP data and assess which of the two models fits LFPs better. This will test evidence for ephaptic coupling. We first discuss the ephaptic extension of the neural field model below.

Above we presented a model of neural activity (cf. Equations (1)) describing current flow within an ensemble. This current generates an electric field in extracellular space, *E^e^*. This can directly influence individual neurons, a phenomenon known as *ephaptic coupling* (Anastassiou et al., 2011; Fröhlich and McCormick, 2010; Rebollo et al., 2021; Ruffini et al., 2020; Schmidt et al., 2021). Ephaptic coupling describes interactions between the brain’s electric fields and neural activity, that is, interactions between *E^e^* and *V^m^*. (Danner et al., 2011) and (Goldwyn et al., 2017) showed that ephaptic effects result in perturbations (small increases) of transmembrane potential by adding the value of the extracellular potential on the membrane *V*_0_^*e*^. They described these increases by replacing *V^m^* ≍ *V^m^* +*V ^e^* in the term capturing local recurrent input as a result of diffusion. In other words, they added an ephaptic current to the diffusion current that changes the transmembrane potential. We did the same here. We replaced *V^m^* ≍ *V^m^*+*V^e^*_0_ in the integral in Equation (1) that describes diffusion of recurrent input in the ensemble and obtained

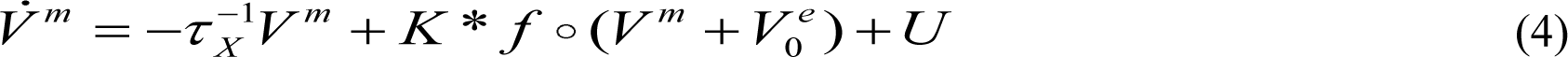

We called this the *ephaptic model*. Compared to Equation (1), Equation (4) suggests that the rate of change of depolarization comprises the same three terms as before and additionally, perturbations due to extracellular potential *V^e^*_o_. The ephaptic model is used twice below. First in *Methods* to derive the mathematical expression of ephaptic coupling. Then in *Results,* to find evidence of ephaptic coupling using Bayesian Model Comparison (BMC).

### A model of the ensemble electric field

We saw above that current flow within the neural ensemble generates an electric field in extracellular space 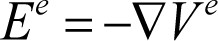, where *V^e^* is the corresponding potential. In (Pinotsis and Miller, 2022), we introduced a model of this electric field based on the *bidomain model* (Goldwyn et al., 2017; Mc Laughlin et al., 2010). Below, we summarize the main points of this model. For more details, the interested reader is invited to consult (Goldwyn et al., 2017; Mc Laughlin et al., 2010; Pinotsis and Miller, 2022).

The bidomain model assumes that dendrites of cortical pyramidal cells comprising neural ensembles extend parallelly. Although they have a complicated geometry, this symmetry allows one to replace the branched dendrites trees by a cylindrical fiber (Rall, 1998). This is the same symmetry as that of the current dipole approximation to cortical sources widely used in human electrophysiology (Hämäläinen et al., 1993; Lindén et al., 2010; Nunez and Srinivasan, 2006).

In this model, pyramidal neurons are aligned to produce an EF parallel to apical dendrites and receive synchronous input. Current flowing in neurons gives rise to dipole sources (Buzsáki et al., 2012; Pesaran et al., 2018). The extracellular space of each pyramidal neuron is described by a cylindrical fiber (small blue cylinders in Figure 1B). Using the principle of superposition from electromagnetism, extracellular spaces can be combined into a unified extracellular space of the neural ensemble. Thus, the individual cylindrical fibers of Figure 1B (for each neuron) are replaced can be replaced by the larger fiber surrounding the ensemble (light blue cylinder in Figure 1C). The boundary between extracellular and intracellular space has the same symmetry and is denoted by a grey cylinder in Figure 1C.

The bidomain model assumes spatial homogeneity and temporal synchrony similarly to the well known dipole approximation. EF model estimates are a bound on realistic values of EF: actual EFs will be smaller when these assumptions fail. Note that this does not change qualitive results, like ephaptic coupling discussed below as the extracellular and intracellular spaces can be split into smaller parts (cylindrical fibers) where symmetry and synchrony still apply.

This electric field *E^e^* is the result of the discontinuity in the electric potential *V^e^*_o_ − *V^i^*_o_ that gives rise to electric dipole sources and transmembrane currents 1/ *r_i_* Δ*V^m^* (*V^e^*_0_ and *V^i^*_0_ are the values of the extracellular and intracellular EPs on the two sides of the membrane). Intuitively, *E^e^* is the potential difference over unit distance. Alternatively, *E^e^* expresses the force to which an ion is subjected to, while in extracellular space, divided by its charge (Jackson, 1999).

Because of symmetry, the extracellular field and potential depend on two spatial variables (*x*, *y*), not three. The variable *x* parameterizes the location on the axis of the cylinder in Figures 1C and 1D and *y* a direction orthogonal to this axis. According to the bidomain model, the extracellular potential *V^e^* at a point *P*(*x*, *y*) in the extracellular space^1^ is given in terms of the Fourier Transform *V̑^m^* of the transmembrane potential *V^m^* by the following expression, see Equation (17) in (Pinotsis and Miller, 2022b) and *Supplementary Material* for more details,

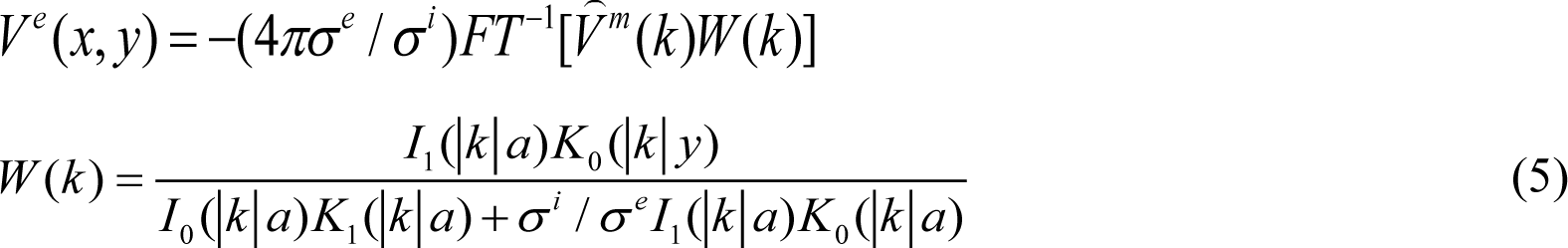

Note that because of cylindrical symmetry the functions appearing in the second line of Equation (4) do not depend on *x*. They just depend on *y* and the ones appearing in the denominator are evaluated for *y* equal to the radius of the grey cylinder, *y=a* (the cylinder separating intracellular and extracellular space, like a membrane). Here, *σ ^l^*, *l* = {*e*,*i*}, are the extra-and intra-cellular space conductivities and *I*_0_ (*y*), *I*_1_ (*y*), *K*_0_(*y*), *K*_1_(*y*) are modified Bessel functions of the first and second kind (Abramowitz et al., 1988). In brief,

### Ephaptic coupling, Synergetics and the stability of the electric field

In the next two sections of the *Methods*, we include some mathematical arguments that motivate hypotheses tested in *Results.* These involve analytical, ie. pen and paper calculations. Above, we summarized a model of the electric field generated by neural ensembles. In (Pinotsis and Miller, 2022b), we used this model to compute the EF corresponding to neural ensembles maintaining different memory representations. We found that EFs were more stable than neural activity and contained relatively more information. We suggested that this stability allows the brain to control the latent variables that give rise to the same memory. In other words, we hypothesized that EFs can sculpt and herd neural activity and can act as “guard rails” that funnel the higher dimensional variable neural activity along stable lower-dimensional routes.

Below we provide further mathematical arguments in support of the above hypothesis: that bioelectric fields guide neural activity. In the *Results* section, we test this hypothesis, using data from a spatial delayed saccade task.

We were interested in interactions between variables expressed at different spatial and temporal scales: bioelectric fields and neural activity. Thus, we used a theory that describes interactions underlying spontaneous pattern formation in biological and physical systems known as *synergetics* (Haken, 1987; Jirsa and Haken, 1996). Synergetics studies how individual parts –in our case neurons— produce structures; here, memory representations. It suggests that a biological system, like a neural ensemble, is constrained by so called *control parameters* that impose limitations. When control parameters change, the structures change. A simple example of a control parameter is temperature. When it changes, the state of water molecules can change from solid, to fluid, to air. In synergetics language, the individual elements of the system, e.g. molecules, are called enslaved parts. This is because they are controlled by temperature. Besides control parameters and enslaved parts, synergetics also considers order parameters, that is, low dimensional descriptions of collective dynamics, like the average transmembrane potential *V^m^* that we studied here or other latent variables (Gallego et al., 2020; Yu et al., 2008) like effective connectivity components (Pinotsis et al., 2017b). A crucial distinction between control and order parameters is how fast they evolve. When there is a perturbation, like new input to a brain area, the order parameters and enslaved parts evolve fast and the control parameters slowly. Control parameters are very stable compared to order parameters. To put it differently, synergetics suggests a temporal hierarchy comprising, slow control parameters, like temperature or energy (Ditzinger and Haken, 1989), faster order parameters and very fast enslaved parts (e.g. oscillations/spiking(Miller et al., 2018)).

Below, we will use the theory of synergetics to provide a mathematical formulation of ephaptic coupling, that is, the interactions between the ensemble electric field, *E^e^*, and the average transmembrane potential *V^m^*. We will present some theoretical arguments that motivate the hypothesis that a slow EF *E^e^* acts as a control parameter that enslaves faster neural activity *V^m^*. In *Results*, we will test this hypothesis and ask whether ephaptic coupling can be detected in *in vivo* neural data.

To describe extracellular field – transmembrane potential *E^e^*-*V^m^* interactions, our starting point is equations that express one quantity in terms of the other, that is *E^e^* in terms of *V^m^* and vice versa. These are Equations (4) and (5): the evolution of transmembrane potential *V^m^* in terms of the extracellular EP *V^e^* is given by the ephaptic model (4). Also, *V^e^* in terms of transmembrane potential *V^m^* is given by the bidomain model (5). To perform pen and paper calculations, we need algebraic equations (ie. equations without the inverse Fourier transform *FT^-1^*). Thus, in Supplementary Methods we show how we one can rewrite Equation (5) as a differential algebraic equation, see Equation (6) below. For simplicity, we assume that the LFP electrode is at a large distance compared to the size of the neural ensemble: the radius *a* of the fiber separating the intra- and extra-cellular spaces (grey cylinder) is very small compared to the vertical distance *y* to the location of the LFP electrode, *a* << *y*, *cf.* squashed grey cylinder in Figure 1D.

From trial to trial the remembered stimulus changes. Thus the EP and the corresponding EF also change, see (Pinotsis and Miller, 2022b) for details. Assuming a fixed point attractor (steady state), Equation (5) can be written as (see *Supplementary Material* for details)

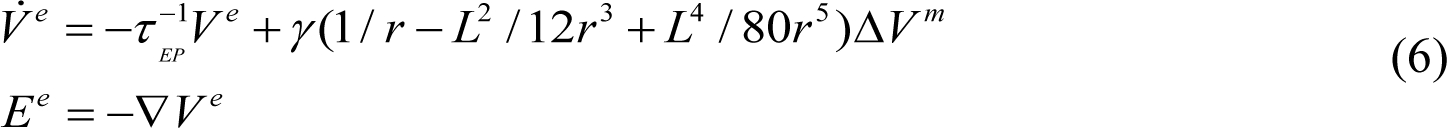

where *τ*^−1^_*EP*_ is the rate with which *V^e^* decays to its resting value *V_S_^e^*.

Equation (6) expresses the dynamics of the extracellular EP *V^e^* in terms of the transmembrane potential *V^m^*. To describe interactions between these potentials and the corresponding electric fields, we then applied the *slaving principle* from synergetics (Haken, 1987). This predicts that control parameters evolve more slowly and constrain order parameters and enslaved parts. Examples of the general slaving principle can be found in physics and biology (Haken, 2012). Haken and colleagues have shown that varying the temperature (control parameter) of a fluid heated from below, various spatial patterns of fluid molecules occur. Also, that attention can be thought of as control variable in multi-stable perception (Basar et al., 2012; Ditzinger and Haken, 1989).

During working memory delay, the *slaving principle leads to ephaptic coupling*: it predicts that extracellular EP, *V^e^*, enslaves neural activity described by the transmembrane potential *V^m^*. To confirm this, consider the following expansion of *V^m^* and *V^e^* in terms of Fourier series 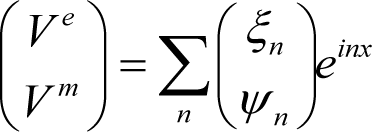. Then, substituting these expansions into Equations (4) and (6), we ç*_V m_* ÷ åç*_ψ_* ÷*^e^* è ø *n* è *n* ø obtain evolution equations for the Fourier coefficients or modes

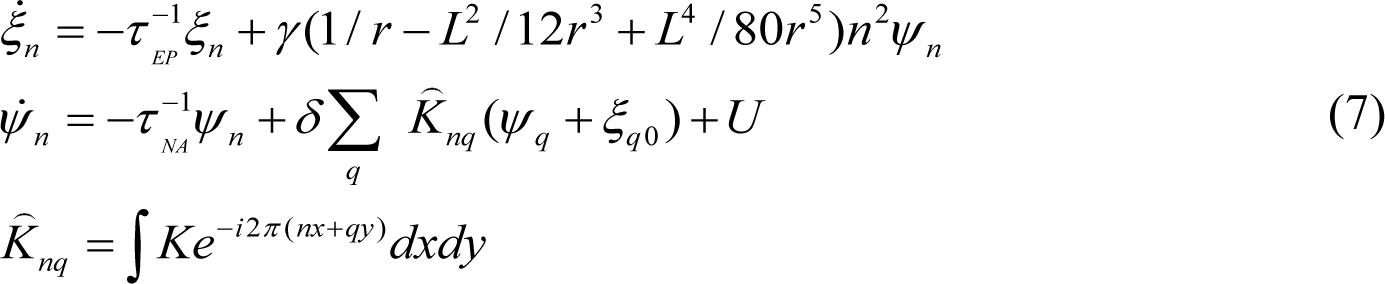

*ξ_n_* and *ψ _n_* are called the Fourier coefficients or modes of the extracellular potential and neural activity. Below, we call them modes. *ξ_q_*_0_ are the values of the extracellular EP on the exterior of the ensemble membrane (surface of grey cylinder in Figure 1B). Intuitively, a Fourier expansion implies that *V^m^* and *V^e^* are superpositions of planar waves *e^inx^* with amplitudes given by *ξ_n_* and *ψ _n_*.

We have replaced Equations (4) and (5) that describe the coupling between the extracellular potential and neural activity, *V^e^* and *V^m^*, by Equations (7) that describe the same coupling in terms of modes. Note that in the second equation (7) that the rate of change of neural activity modes, *ψ _n_*, depends on values of the extracellular potential modes *ξ_q_* _0_ on the exterior of the membrane and exogenous stochastic input *U*.

We can now apply the slaving principle of synergetics. This suggests that in Equations (7), one can distinguish between slow and fast modes. Equations –as usual—provide a formalism and motivate experimental tests, they cannot replace these tests. In (Pinotsis and Miller, 2022b), we found that the electric field was more stable than neural activity. Correlations of single trial estimates of electric fields were higher than correlations of similar neural activity estimates. Based on these findings and evidence from a multitude of ephaptic coupling studies, see e.g. (Anastassiou and Koch, 2015), it is sensible to assume that extracellular potential modes *_ξ_* are slow and the transmembrane potential modes *ψ _n_* are fast. In that case, the damping constant for the extracellular potential would be much smaller than the damping constant for neural activity 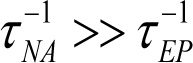 (adiabatic approximation ;Haken, 1987).

Assuming 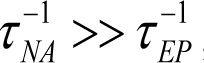, Equations (7) suggest that the instantaneous values of fast relaxing quantities, like the transmembrane potential modes *ψ _n_*, depend on slowly varying quantities, like the extracellular potential coefficients *^ξ^q* 0 above, which slave them (Haken, 1987). Electric fields enslave neural activity. This is ephaptic coupling formulated in the language of synergetics –as a special case of the slaving principle. Equations (7) are then the mathematical expression of ephaptic coupling. In (Haken, 2012, 1987), several Equations similar to (7) are presented in the context of physics and biology and similar coupling between fast and slow quantities is discussed.

Note that Equations (7) are *not* used for calculations in *Results*. Had we used them in our calculations, we would have had to prescribe the rate constants 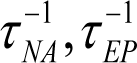 a priori. This would bias our conclusions. Equations (7) are useful because they *motivate* a hypothesis that is tested in *Results* –independently of Equations (7). This hypothesis is that electric field modes *^ξ^q* 0 are slow and neural activity modes *ψ _n_* are fast. A theoretical argument in support of this hypothesis is a common assumption in bio-electromagnetism about the EF being quasi-static: the tissue impedance on top of resistance (or more generally reactance) is assumed to be negligible and electromagnetic propagation effects can be ignored (Nunez, 1998). In other words, the electric field is assumed to be relaxing very slowly compared to quickly relaxing neural activity.

Assuming that neural activity is enslaved by the electric field has another implication. It suggests that instantaneous values of neural activity are given in terms of instantaneous values of the slower fields. During the delay period of the memory task considered here, one can assume fixed point dynamics. In other words, the transmembrane potential can be assumed to be in equilibrium, thus *ψ _n_* = 0. Then, Equations (7) yield these instantaneous values of neural activity determined by emerging fields. One can express *ψ _n_* in terms of *ξ_n_*

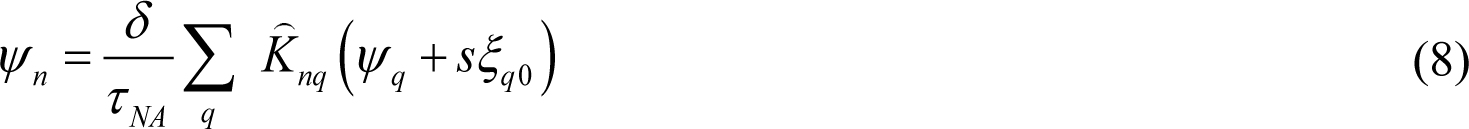

This equation describes how the fast modes of neural activity are enslaved (driven) by the slow, *stable* modes of the electric field.

To sum, the slaving principle from synergetics predicts that stable electric fields enslave neural activity. Mathematically, this result is expressed via Equations (7) and in neuroscience it is called ephaptic coupling. The slaving principle distinguishes between stable and unstable quantities, like the modes *ξ_n_* and *ψ _n_*. It suggests that the evolution of fast unstable modes is determined by stable modes. The latter determine the instantaneous values of the former: here the electric field determines neural activity, see Equation (8). This is also related to critical slowing where some modes are strongly correlated over time, e.g. (Bassett and Bullmore, 2009; Chialvo, 2010; Kitzbichler et al., 2009), see also (Pinotsis and Miller, 2022b).

### Ephaptic coupling across engram complexes

The distinction between stable and unstable modes can be obtained using a mathematical theory known as linear stability analysis. Linear stability analysis of neural network models is often used to express brain responses in terms of key anatomical and biophysical parameters, e.g. (Coombes, 2005; Jirsa and Haken, 1996; D.A. Pinotsis et al., 2013; Pinotsis and Friston, 2010). It can also be extended to include nonlinear terms, see (Basar et al., 2012; Haken, 1987). Here, we use linear stability analysis to motivate a hypothesis about engram storage in memory networks that will be tested in *Results*: that ephaptic coupling occurs across engram complexes. *If ephaptic coupling occurs in a brain area and this exchanges memory information with other brain areas then ephaptic coupling will occur in those areas too.* Below, we present mathematical argument in support of this hypothesis for two areas. Generalization to an arbitrary number of areas can be done by induction.

Consider two neural ensembles in brain areas (1) FEF and (2) SEF. Dynamics of ensemble activity are given by a system of neural fields of the form of Equation (4). Similarly to Equation (4) above, ephaptic coupling suggests *V ^jm^* depends on EP *V^je^*_0_ (its boundary value at the membrane exterior) via the following expressions:

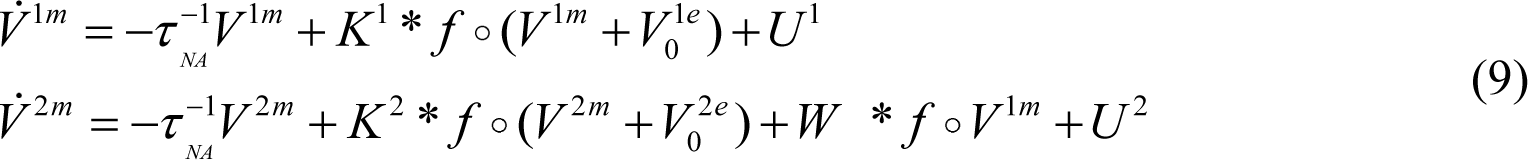

Here, *W* is the feedforward connectivity matrix whose entries are weights that scale downstream input to SEF from FEF (Grossberg, 1967; Wilson and Cowan, 1973). Note that in *Results* we did *not* use predicted data from this model (9). This is just used here for the sake of mathematical analysis. In the linear stability regime, we can assume that the transmembrane potential *V ^im^* of each ensemble (identified by the upper index *j =*1,2) includes perturbations in the form of planar waves around baseline *V ^io^*, that is an equation of the form *V^jm^* ∼ *V^jo^* + *e^βt+ikx^* (Grindrod and Pinotsis, 2011; Pinotsis and Friston, 2010). For mathematical convenience, we consider a vector of extracellular and transmembrane potential functions for the two areas

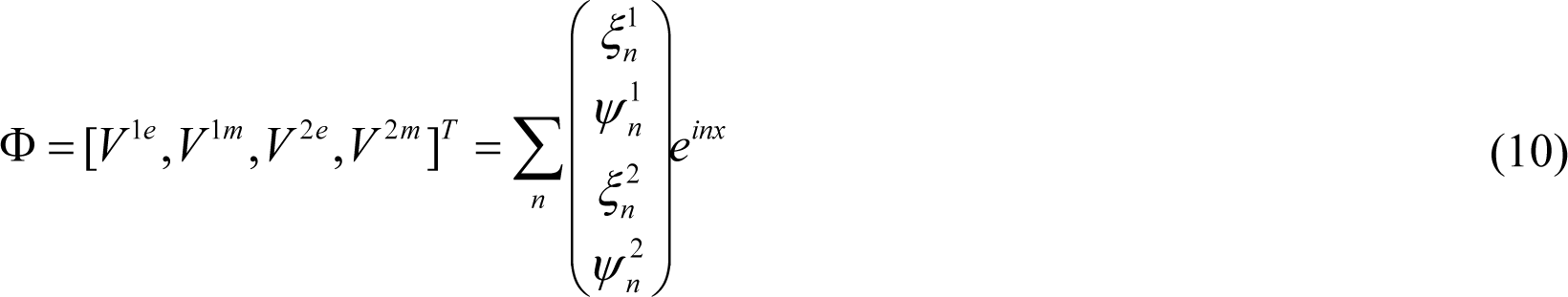

Upper indices denote the area and lower indices the mode order. In the previous section we saw that the slaving principle suggests that the slow, stable field modes ξ^1^_*n*_ and ξ^2^_*n*_ will constrain *ψ*^1^_*n*_ and *ψ*^2^_*n*_. The order of the expansion (10), *n*, (how many modes are needed to faithfully represent the dynamics) can be found using a model fitting procedure (e.g. maximum likelihood or similar) using real data. We will consider this elsewhere. Since we here focus on mathematical arguments, for simplicity, we assume that the first two modes explain most of the observed variance, that is, we keep terms up to 2^nd^ order in Equation (10) (*n=*1,2)

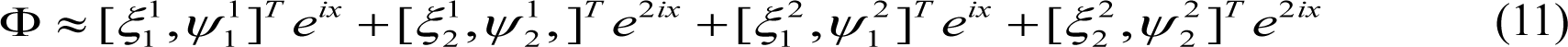

Substituting the above expression in Equations (9) and using the first of Equations (7), we obtain a system of equations

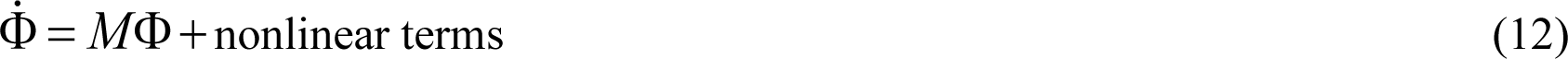

where the matrix *M* can be expressed in terms of 4×4 matrices *A,B,C* and *D*, 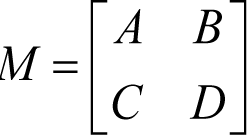 defined in the *Supplementary Material.* Further, the matrix *D* can be written as 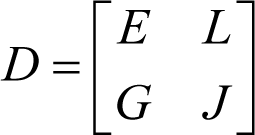 in terms of *2×* matrices *E,L,G* and *J* also included in the Supplementary Material. Equation (12) is a linearized system that describes the coupling of extracellular and transmembrane potentials in the two areas in terms of connectivity. Mathematically, for the system to have a solution, that is, for the modes in all areas to exist, the determinant of the matrix *M* needs to be different than zero, det(*M*) ≠ 0. Existence of a solution of a linear system when the determinant of the coefficient matrix is non zero is a standard result in Linear Algebra (Strang, 2006). Note that *M* is called the coefficient matrix of the linearized system given by Equation (12).

But what does existence of solution mean? Intuitively, it means that one can find functions that satisfy these equations. This, in turn, implies that there are some functions ψ_*j*_^1^, ξ_*j*_^1^ and ψ_*j*_^2^, ξ_*j*_^2^, that is, some extracellular and transmembrane potentials that can describe electrical activity in FEF and SEF. The implied assumption here is that there is also a feedforward connectivity matrix *W* (recall Equation (9)) so that FEF and SEF form an engram: there is input from one area to the other. To sum, Equation (12) and the condition det(*M*) ≠ 0 are just a mathematical expression of the simple fact that ensembles FEF and SEF are connected and generate some activity and electric fields. By applying the identity det(*M*) = det(*A* − *BD*^−1^*C*)det(*D*) (Abramowitz et al., 1988), we obtain

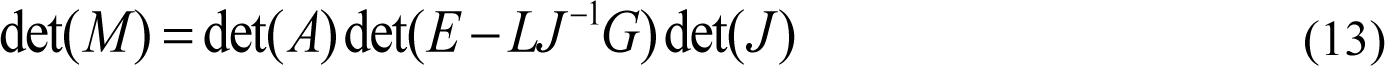

Thus, the condition det(*M*) ≠ 0, requires that det(*J*) ≠ 0 and det(*A*) ≠ 0 ; the determinants of matrices *J* and *A* should also be non-zero. *J* is defined by

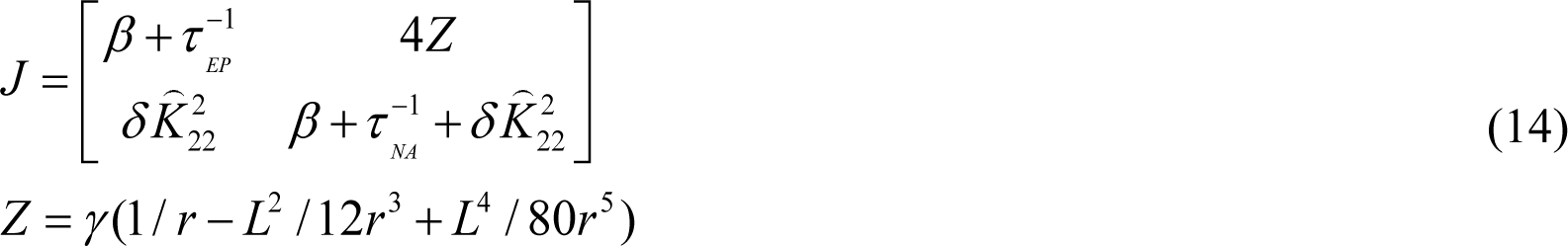

In other words, *J* is the matrix of coefficients in a linearized system of equations describing the coupling between the second extracellular and membrane potential modes in the second region:

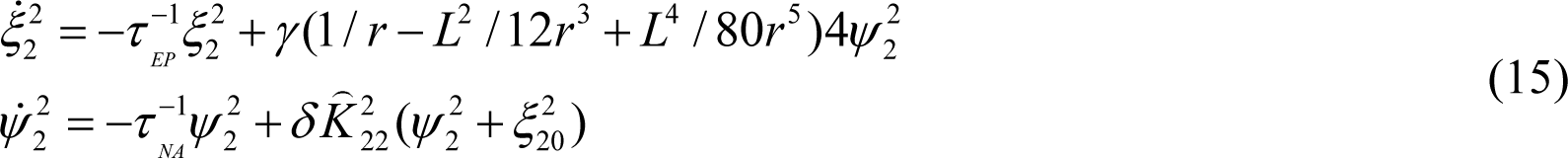

Then, the condition det(*J*) ≠ 0 implies that the above system has a solution, i.e. there are some functions (modes) that describe the extracellular and transmembrane potentials. det(*J*) ≠ 0 also includes some additional piece of information. This is due to the similarity of these Equations with Equations (7). In the previous section, we found using synergetics that Equations (7) are the mathematical expression of ephaptic coupling. Equations (15) are the same as (7) when we consider the second order modes (denoted by the lower index “2” in *ξ*^2^) in the second area (SEF; (denoted by the upper index “2”). Thus, Equations (15) are the mathematical expression of ephaptic coupling for second order modes in the second area. In other words, if we assume that Equations (15) hold (mathematically, a solution exists) the transmembrane and extracellular potential modes will be linked via ephaptic coupling. Similarly, the condition det(*A*) ≠ 0 means that the modes in the first area will also be linked via ephaptic coupling.

We turn to Equation, (13). This says that if the determinants of the matrices *M* and *J* are non-zero, then the determinant of matrix *A* will also be non-zero (the same is true if we replace *J* by *A*). In symbols, if det(*M*) ≠ 0 and det(*J*) ≠ 0 then det(*A*) ≠ 0. Above we saw what each of these three conditions means. Following these earlier interpretations, we can put the last mathematical statement into words: Assuming that FEF and SEF are connected and generate activity and electric fields (det(*M*) ≠ 0) and that there is ephaptic coupling in the second area (det(*J*) ≠ 0) there will be ephaptic coupling in the first area (det(*A*) ≠ 0 ; or the other way around where we replace *J* by *A*). In brief, assuming that ephaptic coupling occurs in one area, then ephaptic coupling will also occur in the other area. By induction, we can show the same result for an arbitrary number of areas that form a memory network or engram complex.

### Granger causality

To test for information transfer between different spatial scales (emerging electric fields and neural activity) and brain areas (FEF and SEF), we used Granger causality (GC; (Geweke, 1982; Granger, 1969)). GC quantifies how the history (past samples) of variable A improve prediction of unknown samples (future samples) of a different variable B. It is based on generalized variances or log likelihood ratios that quantify whether a regression model including variable A fits future samples of variable B better than the restricted regression model based on variable B samples only (Friston et al., 2013). Following (Barnett and Seth, 2014),we evaluated GC as follows: we first used model based VAR modeling to calculate regression coefficients from our data, similar to a discrete stationary vector stochastic process. First, one determines an appropriate order of a VAR model using an information criterion or cross validation(Burnham and Anderson, 1998). Then, a log-likelihood ratio*^F^A*→*B* of residual covariance matrices is computed. This corresponds to the full and restricted VAR models and quantifies the *GC strength*, that is, whether the prediction of future values of the variable *B* improves significantly after including past values of *A.* This can be computed using Granger’s *F-*test for univariate problems or a chi-square test for a large number of variables (Geweke, 1982; Granger, 1969). GC is often used for the analysis of time series (samples are obtained using measurements at different moments in time). Here, we used GC after considering spatial samples, that is, we obtained measurements at different locations in the neural ensemble and extracellular space. This is discussed further in *Results*.

### Representation Similarity Analysis

We used Representation Similarity Analysis(Kriegeskorte et al., 2008) (RSA) to assess the similarity of information representation across different brain areas. RSA uses Dissimilarity Matrices (DMs) to summarize how stimulus information is represented by brain responses. Following (Pinotsis et al., 2019), we built DMs based on time correlations that are thought to underlie working memory representations (Inagaki et al., 2017; Wallis et al., 2015). Each DM entry contained the dissimilarity between trials corresponding to different remembered cued locations. Thus, DMs describe pairwise differences in patterns of neural activity corresponding to different stimuli. To understand whether similar information (cued location) was encoded in different brain areas, we computed the dissimilarity between brain DMs.

Following (Kriegeskorte et al., 2008), the dissimilarity between dissimilarity matrices, known as deviation, was the correlation distance (1-Spearman correlation; Spearman was used as it does not require a linear correspondence between these matrices contrary to Pearson correlation). Deviations between DMs quantify matches between representation content of brain responses (Kriegeskorte, 2011). They measure the correlation distance between each DM and quantify differences of differences: How different are the corresponding pairwise differences in neural activity or electric fields. After calculating deviations of DM matrices one can assess significant correspondence between information stored in different brain areas (Diedrichsen and Kriegeskorte, 2017; Peterson et al., 2018).

## Results

### Ephaptic coupling in in—vivo memory delay data

We first asked whether we could find evidence for ephaptic coupling in our data. We examined in vivo LFPs acquired from FEF and SEF during delay in a spatial WM task (Jia et al., 2017; Pinotsis et al., 2017b). In (Pinotsis and Miller, 2022b), we analysed the same data from FEF only. Here we extended our analyses to the FEF-SEF memory network (engram complex).

To assess evidence of ephaptic coupling in our data we used computational modeling. We considered two variants of the same model: with and without ephaptic effects (ephaptic and non ephaptic). First, we fitted the models to LFP data and compared their fits. Second, we used model predictions and Granger causality to assess evidence of ephaptic effects. This is discussed in the next section. Below, we discuss computational models and their fits.

In earlier work (Pinotsis et al., 2017b), we obtained predictions of the activity of neural ensembles maintaining different cued locations. Transmembrane depolarization was described by a neural field model trained as an autoencoder, that we called a deep neural field. The term “deep” reflects the bottleneck architecture of the Restricted Maximum-Likelihood (ReML) algorithm used for training. The model was trained using the same LFP data as those considered in the analyses below, see Methods.

Here, we obtained new predictions of the activity of neural ensembles by extending the model of (Pinotsis et al., 2017b) to include ephaptic coupling (Methods), see also (Goldwyn et al., 2017). In other words, our analyses below used two sets of predictions of neural activity: with and without ephaptic coupling. Predictions without ephaptic coupling were obtained in (Pinotsis et al., 2017b). Predictions of neural activity with ephaptic coupling were obtained here following (Danner et al., 2011; Goldwyn et al., 2017). These were obtained in two steps: First, we calculated the extracellular electric potential generated by the neural ensemble using a model from bioelectromagnetism (bidomain model) introduced in (Pinotsis and Miller, 2022), see also (Goldwyn et al., 2017; Mc Laughlin et al., 2010). Second, we added an ephaptic current to the local recurrent input to the neural ensemble that changed its activity. The ephaptic current was an additional current resulting from effects of the extracellular electric potential near the ensemble.

To look for evidence of ephaptic coupling, we fitted the predictions of the ephaptic and non-ephaptic models to LFP data and evaluated goodness of their fits. Model parameters were the same as in (Pinotsis et al., 2017; Pinotsis and Miller 2022b). These are included in the Supplementary Table. We used Bayesian Model Comparison (Friston et al., 2007; K. Friston, 2008; Pinotsis et al., 2014) to find the model that fit the data best (Pinotsis et al., 2018). The ReML algorithm was used for fitting (K. Friston, 2008). We also used previously unseen data (data not used for training) to avoid data leakage.

We compared the evidence of the two models (how well a model could explain the data), the ephaptic and the non-ephaptic. If the fit of the ephaptic model was better, this would provide evidence of ephaptic coupling under the assumption that models are plausible. The validity of the original neural field model (without ephaptic coupling) has been assessed previously: variance explained by the model was about 40%, see Supplementary Figure 3A and (Pinotsis et al., 2017). In (Pinotsis and Miller, 2022) we also showed that the model obtained the same neural ensemble connectivity as that obtained using two independent methods: k-means clustering (Humphries, 2011) and high dimensional SVD (Carroll and Chang, 1970; Williams et al., 2018). The above results support the validity of the original model (non ephaptic). We will return to the validity of the extended model (ephaptic model), after we discuss the results of model comparison below.

To compare models and evaluate their fits, we used model evidence. This was computed using a Free Energy approximation. Free Energy is a cost function borrowed from autoencoders that we used to measure goodness of fit. Inference used single trial data and the principal axes as input to infer connectivity, similar to Dynamic Causal Modeling (DCM) and other model fitting approaches (Freestone et al., 2014; Oesterle et al., 2020; D. A. Pinotsis et al., 2012). Having obtained the Free Energy, one can computer the Bayes factor (BF; Kass and Raftery, 1995). BF> 3 suggests that the model with the higher Free Energy explains the data better. BF can be thought of as a probabilistic analogue of the odds ratio used in frequentist statistics. This corresponds to a posterior probability of 95% for the winning model. Here, BF describes how likely is the ephaptic model to have generated the sampled LFPs compared to the non ephaptic model, under a fixed effects assumption (same model for all trials).

BF results are shown in Figure 2A (vertical axis). These are averaged over trials for each cued location. The horizontal axis shows the 6 different locations (angles) cued to hold in working memory. Blue bars denote the BF after fitting FEF data, while red bars after fitting SEF data. A positive BF implies that the non-ephaptic model was more likely; a negative BF that the ephaptic model was. The arrow at the right hand side of Figure 2A facing upwards includes the letters NE = non-ephaptic, while the downwards facing arrow, the letter E=ephaptic wins. BF bars pointing “downwards” provides evidence of ephaptic coupling.

**Figure 2.**
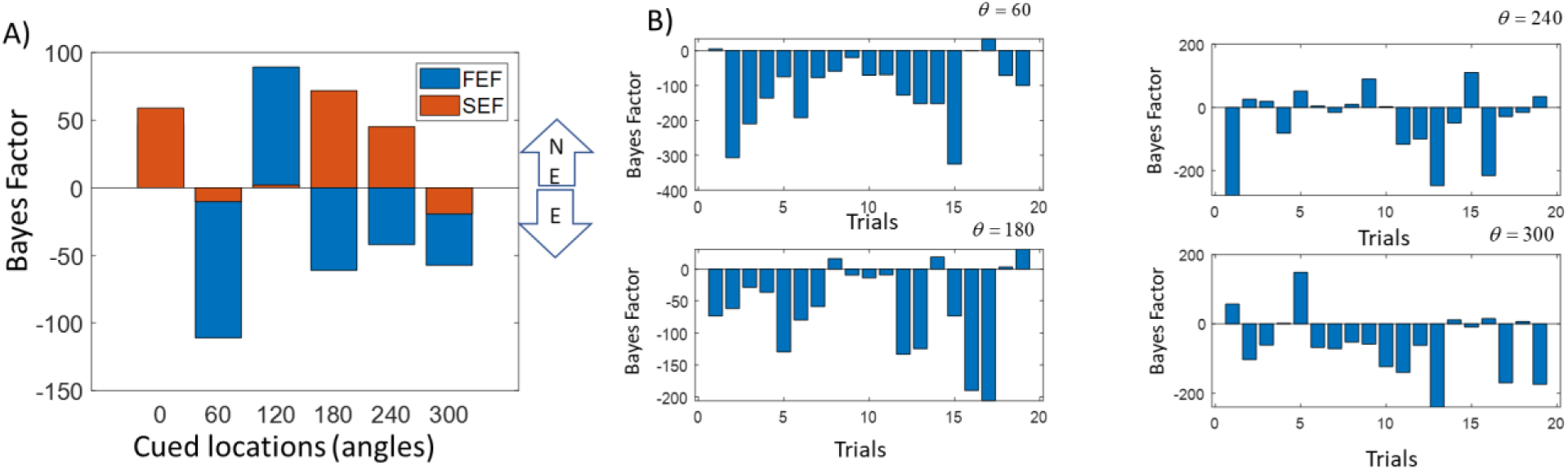
(A) Bayes factor (BF) for different cued locations (horizontal axis). Blue bars denote the BF after fitting FEF data, while red bars after fitting SEF data. A positive BF implies that the non-ephaptic model was more likely; a negative BF that the ephaptic model was. BF bars pointing “downwards” provides evidence of ephaptic coupling, denoted by the E inside the lower arrow. NE in the upper arrow stands for “non-ephaptic”. (B)Bayes factor for individual trials and specific cued angles when the ephaptic model wins. Different trials are shown on the horizontal axis. The corresponding cued angles are shown at the top right corner of each plot. The ephaptic model fits the data better for most trials.

Using model comparison, we found that in FEF, the ephaptic model was more likely for cued locations at *θ* = 60,180, 240 and 300 degrees (BF=*-120,70,45* and *55* respectively; blue bars in Figure 2A). To make sure the ephaptic model fitted single trial data better, Figure 2B shows the BF for individual trials for *θ* = 60,180, 240 and 300 degrees, i.e. when the ephaptic model was more likely in FEF (results for other angles are similar, not shown). Average BF estimates reported in Figure 2A are not driven by outliers. We confirmed that the ephaptic model was better in most trials. BF estimates are between BF=*20-310* for *θ* = 60 degrees, BF=*10-200* for *θ* = 180 and *θ* = 240 degrees, and BF=*5-220* for *θ* = 300 degrees. In SEF (orange bars in Figure 2A), the ephaptic model was more likely for*θ* = 60 and 300 degrees (BF=*-10* and *20* respectively)^2^. Although results were robust over trials, we did not find evidence of ephaptic coupling across all cued locations. Thus, we then asked why there is evidence in favour of ephaptic coupling for some cued locations and not others. Either there was no ephaptic coupling in these cases or the model overfitted. The first explanation is refuted by the results of the next section that assesses evidence of ephaptic coupling using a different method, Granger causality. The second explanation is consistent with these results and also follows from a careful consideration of model predictions—that also reveals limitations of the non ephaptic model.

We saw above that the original model was found to predict neural activity and connectivity when tested against the LFP data and other methods^3^. The ephaptic model includes small perturbations of transmembrane potentials due to extracellular field effects. We thus focused on these perturbations that we call ephaptic effects (on neural activity). To find them, we subtracted the predictions of the non-ephaptic model from the corresponding predictions of the ephaptic model (averaged over trials). The models predict fluctuations of neural activity around baseline because of endogenous noise driving the neural ensemble in the form of transient non-Turing patterns (patterns that decay back to baseline).

Ephaptic effects are included in Supplementary Figure 1. Supplementary Figure 1A (left) shows the relative percent changes due to the ephaptic coupling for FEF. Similarly, Supplementary Figure 1B (right) shows the corresponding relative changes for SEF. There are six panels in each Figure, each corresponding to a different cued location (angle). This is shown in bottom right of each panel, e.g. the top left panel corresponds to cued location *θ* = 0 degrees. The vertical panel axes show the relative change in principal axis strength with respect to the original principal axis, after including ephaptic coupling. The horizontal panel axes show time in *ms*. Ephaptic effects (amplitudes of neural activity) are expressed as relative increases in amplitude with respect to fluctuations when ephaptic coupling is not considered (i.e. predictions of the non ephaptic, original model). A positive relative change of *α%* implies that the amplitude of neural activity is *α%* larger (or smaller if the change is negative).

Comparing Figures 2A and Supplementary Figure 1, we concluded that the ephaptic model explained the FEF data better only when ephaptic effects were small, i.e. below 40% and cued locations at *θ* = 60,180, 240 degrees. Effects for the case of the remaining two cued locations for *θ* = 0 and 120 degrees are up to 200% (two times larger). Similarly, the model explained the SEF data better for cued locations at *θ* = 0, 60 and 300 degrees when effects were small, i.e. below 6%. For the remaining cued locations at *θ* =120,180 and 240 degrees, ephaptic effects were up to 600% (6 times larger). This suggests that the ephaptic model overfitted large fluctuations –which can be explained from the linearity assumption (Taylor expansion) inherent in its derivation (see *Supplementary Material* and (Pinotsis et al., 2017)).

Below, we did not use the ephaptic model any further. This was only used again in *Methods* to carry out a pen and paper i.e. analytical, derivation of Equations (7) and (12). It was used to formulate mathematical arguments in support of hypotheses tested in *Results* (see *Methods)*. Below, we only used the original, non ephaptic model and Granger Causality. Granger Causality allowed us to test for *nonlinear* interactions between the electric fields and neural activity. This was a second way to assess if there is evidence of in vivo ephaptic coupling in our LFP data (the first was model comparison above). Crucially, Granger Causality also allowed us to obtain the directionality of these interactions. Comparing models above, did not directly assess directionality. We turn to Granger Causality analysis below.

### Top down information transfer from emerging electric fields to neuronal ensembles

Above, we found that, when endogenous fluctuations were small (fractions of fluctuations of membrane potential around baseline), a model in which neural ensemble activity is coupled to the electric field (ephaptic model) explained the LFP data better than a model without ephaptic coupling. We next tested for ephaptic coupling more generally, during large endogenous fluctuations. To do so, we used predictions neural activity from the non ephaptic model considered earlier and Granger Causality (see Equation (1) in *Methods* and relevant discussion). Granger Causality (GC) is a data-driven method for determining the directionality of information flow between stochastic variables (Granger, 1969). Crucially, GC also provides the directionality of the interactions between the electric field and neural activity. In other words, GC allows us to test whether the electric field guides neural activity or the other way around. In (Pinotsis and Miller, 2022a), we suggested that electric fields can act as “guard rails” that funnel the higher dimensional variable neural activity along stable lower-dimensional routes. We tested this hypothesis directly using GC.

Besides the non ephaptic model that gave us predictions of neural activity we also used a model of the electric field, known as the bidomain model, see Equation (5) and relevant discussion in *Methods* and *Supplementary Material* for details. Model parameters for both models are included in Supplementary Table. This model provides predictions of the electric field generated by neural ensembles maintaining different cued locations. Taken together, the non ephaptic and the bidomain model provide two time series, one for predictions of neural activity and another for electric fields. We used the non ephaptic model to get neural activity because its predictions were shown to explain a large part of data variance and to correlate with other methods (see previous section and Footnote 3). Also, the model does *not a priori* assume ephaptic coupling, to avoid biasing results. As with any model, it is just an approximation of the brain’s biology and of the observed neural activity—and similarly for the electric field model. Possible interactions between their predictions would suggest that such interactions could occur in the brain too. Also, the use of GC allowed us to assess nonlinear interactions not considered in the Bayesian model comparison above. Note that we did not use Equations (7) for our results below (because they include rate constants *τ* ^−1^, *τ* ^−1^ and prescribing them a priori would bias our conclusions, see discussion in *Methods*).

Having obtained two time series for neural activity and electric fields, we can assess causal interactions using GC. In its common use, GC is applied to time series data and assesses whether knowing the past of one variable (A) helps predict the future of another variable (B) better than just using the past of B alone. If so, one concludes that information flows from variable A to B. Flow is thought to occur over time, similarly to the flow of a water molecule that flows in a river. In neuroscience, GC is used to describe how information flows in the brain, using sampled time series from different areas(Barnett and Seth, 2014).

One way to compute GC is by first calculating the covariance function, that is, how strongly a time series is related to itself or another time series. This requires *p* samples, that is, measurements at *p* time steps earlier or later (Friston et al., 2013). Implicit in this calculation, there is an assumption of finite *p,* or, that the information flows at a *finite speed* from the variable A to B.

Here, we focused on the information flow between the electric field and neural activity (i.e., electric field and neural activity are the variables A and B). It is well known that interactions involving the electric field transfer information very close to the speed of light, which is practically *infinite*. Thus, the assumption of finite velocity inherent in GC analyses does not hold here. See also the discussion in *Methods* and Equation (8). Following the slaving principle in biology (Haken, 2006, 1987), instantaneous values of neural activity are determined by electric fields, and interactions happen at the speed of light.

Thus, our analysis should be able to deal with field effects being transmitted with practically infinite velocity. This is similar to applications in geophysics where GC and recordings of the earth’s gravitational field are used to e.g. find what kind of minerals exist deep below the surface (Marques et al., 2019). Here, the emerging electric field contains instantaneous information about neural activity in the same way that the gravitational field contains instantaneous information about the masses of minerals underneath. We used this idea from geophysics after replacing the gravitational with the electric field and mineral masses with neural activity.

Because of the practically infinite speed of information propagation (there can be no past or future in time series data of electric fields), we followed a slightly unusual GC analysis where we replaced time with space samples. We considered snapshots of time series and computed the GC over space. We used data from a single time point. Data included the spatial profiles of neural activity and contemporaneous electric field snapshots. We picked the beginning and end of a cortical patch and starting from the beginning, we included all past locations (similar to classical GC where past time points are used) and asked whether knowing the electric field helps predict the value of neural activity in a neighboring (“future” or unknown) location, where activity had not been measured yet, better than using recordings of neural activity alone. GC measures interactions in both directions, thus our analyses answered the reverse question too: whether knowing neural activity helps predict the electric field. Our analyses are summarized in Figure 4.

Following (Barnett and Seth, 2014), we used an *F-*test to assess CG strength *(Methods)*. First, using LFPs from FEF we calculated GC strength and averaged across all time points. Results are shown in Figure 3A for *θ* = 120 degrees. The top right quadrant (from field to activity) has a GC strength of GC=*7.83*, while the bottom left (from activity to field) has a GC strength of GC=*0.04*. Results for other angles are very similar (not shown).

**Figure 3.**
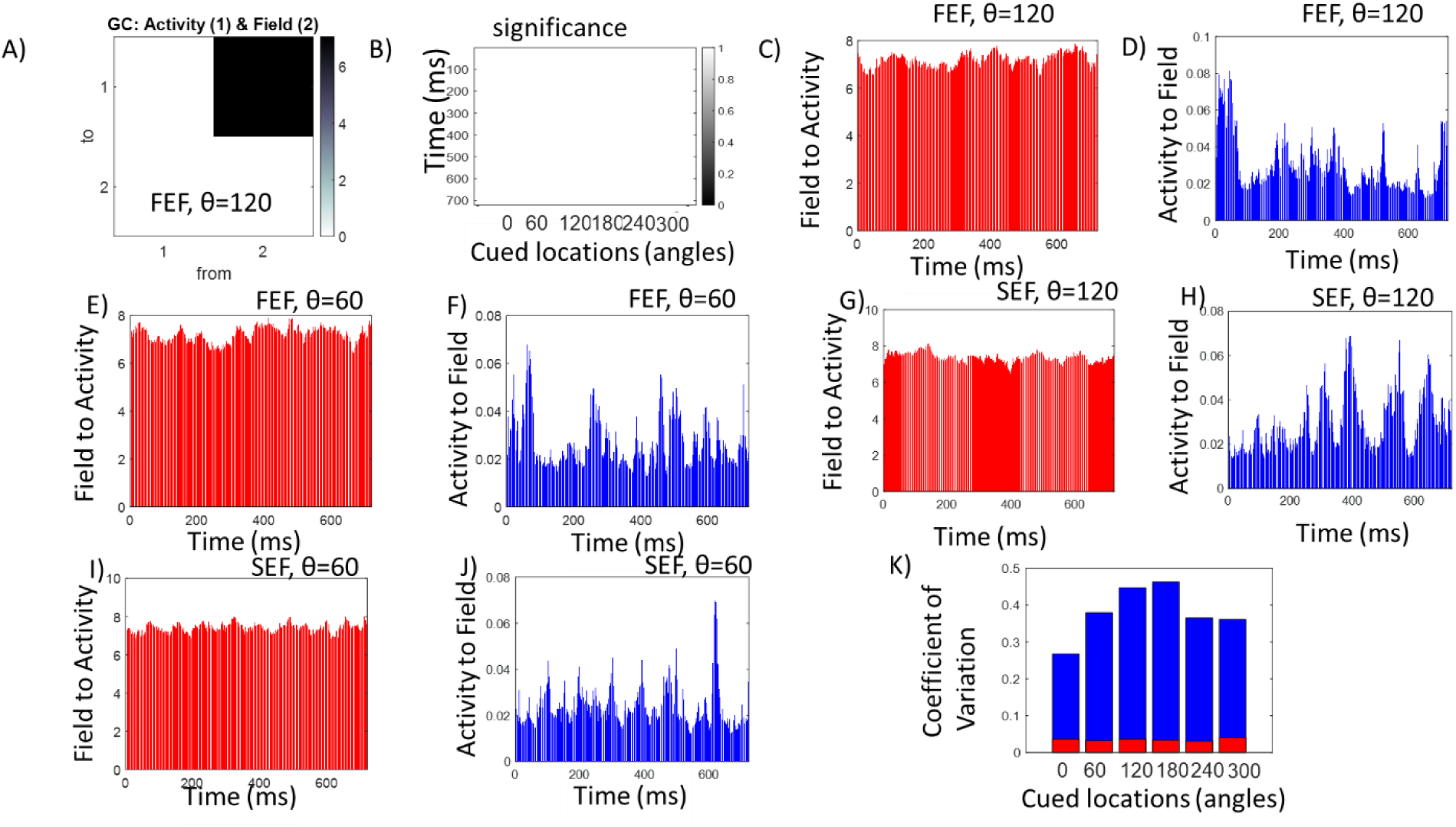
(A) Granger causality (GC) strengths for field-to-neural activity interactions. (B)Significance of GC strengths. All GC strengths were significant (shown as white) across all time points and cued locations that were maintained. (C—F) Examples of individual GC strengths corresponding to each time point during delay for cued locations at θ = 120 and θ = 60 degrees, computed using FEF data. (G – J) Similar to C—F above, for SEF data. (K) Coefficients of variation for GC strengths (vertical axis) for all remembered cued locations (horizontal axis) computed using FEF data. Red bars depict variability in field-to-activity GC strengths and blue bars depict variability in activity-to-field GC strengths.

Figure 3B shows *F-*test significance in the field to activity direction for all cued locations. Time points are shown on the vertical axis and cued locations on the horizontal. All entries are white, (i.e. equal to 1, depicting a logical variable, significant=true) which means that the corresponding GC strength across all time points for *θ* = 120 shown in Figure 3A (after averaging over time) but also all other cued locations was significant. In other words, there were significant field-to-activity GC was significant across all time points and for all remembered angles. Examples of individual GC strengths corresponding to each time point during delay for *θ* = 120 and *θ* = 60 degrees, are shown in Figures 3C – F for FEF and Figures 3G – J for SEF. GC strengths are shown on the vertical axis and time points on the horizontal. Field to activity GC strengths are shown in red, while activity to field GC strengths in blue. In FEF, field to activity GC strengths range between GC=*6.42–7.86* (*θ* = 120 degrees) and GC=*6.41–7.98* (*θ* = 60 degrees). Activity to field GC strengths range between GC=*0.01–0.08* (*θ* = 120 degrees) and GC=*0.01–0.06* (*θ* = 60 degrees). Results for SEF were very similar: field to activity GC strengths range between GC=*6.45–8.19* (*θ* = 120 degrees) and GC=*6.76–8.04* (*θ* = 60 degrees). Activity to field GC strengths range between GC=*0.01–0.07* (*θ* = 120 degrees) and GC=*0.01–0.08* (*θ* = 60 degrees).

All in all, the above results suggest that across all remembered cued locations, GC was much larger in the field to activity than the reverse direction in both FEF and SEF. This confirms our earlier results about in vivo ephaptic coupling in memory ensembles using Bayesian Model Comparison and extends them for all stimuli. The electric field drives the neural activity. It funnels the high dimensional varying neural activity along stable lower dimensional routes – as suggested in (Pinotsis and Miller, 2022a).

Another result from (Pinotsis and Miller, 2022a) was that electric fields were more stable than neural activity. This was confirmed here using GC analysis. Comparing red and blue bars in Figures 3C-J (both FEF and SEF results), we observed that activity-to-field GC strengths varied more over time than field-to-activity GC strengths. This difference in temporal variability between electric field and neural activity is formally assessed using coefficients of variation (CV). Figure 3K shows the CVs for GC strengths (vertical axis) for all remembered cued locations (horizontal axis) using FEF recordings. Red bars depict variability in field-to-activity GC strengths and blue bars depict variability in activity-to-field GC strengths. We found that variability was much higher in the activity-to-field direction. Blue bars corresponding to different cued locations were much larger (CV=*28–47%*) than red bars (CV=*2–4%*). Results for SEF were similar (not shown).

Ephaptic coupling and the stability of the electric field found here (using coefficients of variation based on Granger Causality) follows also from the theory of synergetics. The theory suggests that order parameters, like the electric field, affect enslaved parts, like neural activity (ephaptic coupling). Synergetics also suggests that control parameters (fields) are also more stable than enslaved parts (neural activity). See *Methods* for some mathematical arguments in support of this result.

### Electric fields guide information transfer in engram complexes

Next, we considered causal interactions between electric fields and neural activity in engram complexes across cortical areas. Recall that such complexes include brain areas that maintain memories (Tonegawa et al., 2015a) connected via mono- or poly-synaptic connections. We examined engrams formed by FEF and SEF in our spatial delay task. We studied information transfer between these brain areas using GC. The analyses below are like those in the previous section. The difference is that below, variables A and B are electric fields (or neural activity) from *different* brain areas, as opposed to the same areas considered above. We used predictions of neural activity and electric fields obtained by the non ephaptic and the bidomain models and assessed interactions between them, as before.

We first computed the GC strength based on electric fields in FEF and SEF. This is shown in Figure 4. The corresponding results based on neural activity are shown in Supplementary Figure 2. We first considered at which exact time points interactions between the two areas were significant. These time points are shown in Figures 4A and 4B. Significant interactions are shown in yellow for all cued locations in the FEF to SEF direction in Figure 4A. Remembered cued locations are shown on the horizontal axis while time points on the vertical. Figure 4B has the same format as 4A and shows the corresponding results in the opposite, SEF to FEF direction. For example, for *θ* = 0 degrees, significant electric field interactions in the FEF to SEF directions were observed at sparse intervals between times *t= 290–310ms* and around *t=690ms* (yellow lines in the first column of Figure 4A). In the SEF to FEF direction, such interactions were found around *t=310, 540,490, 540* and *680ms* (Figure 4B).

**Figure 4.**
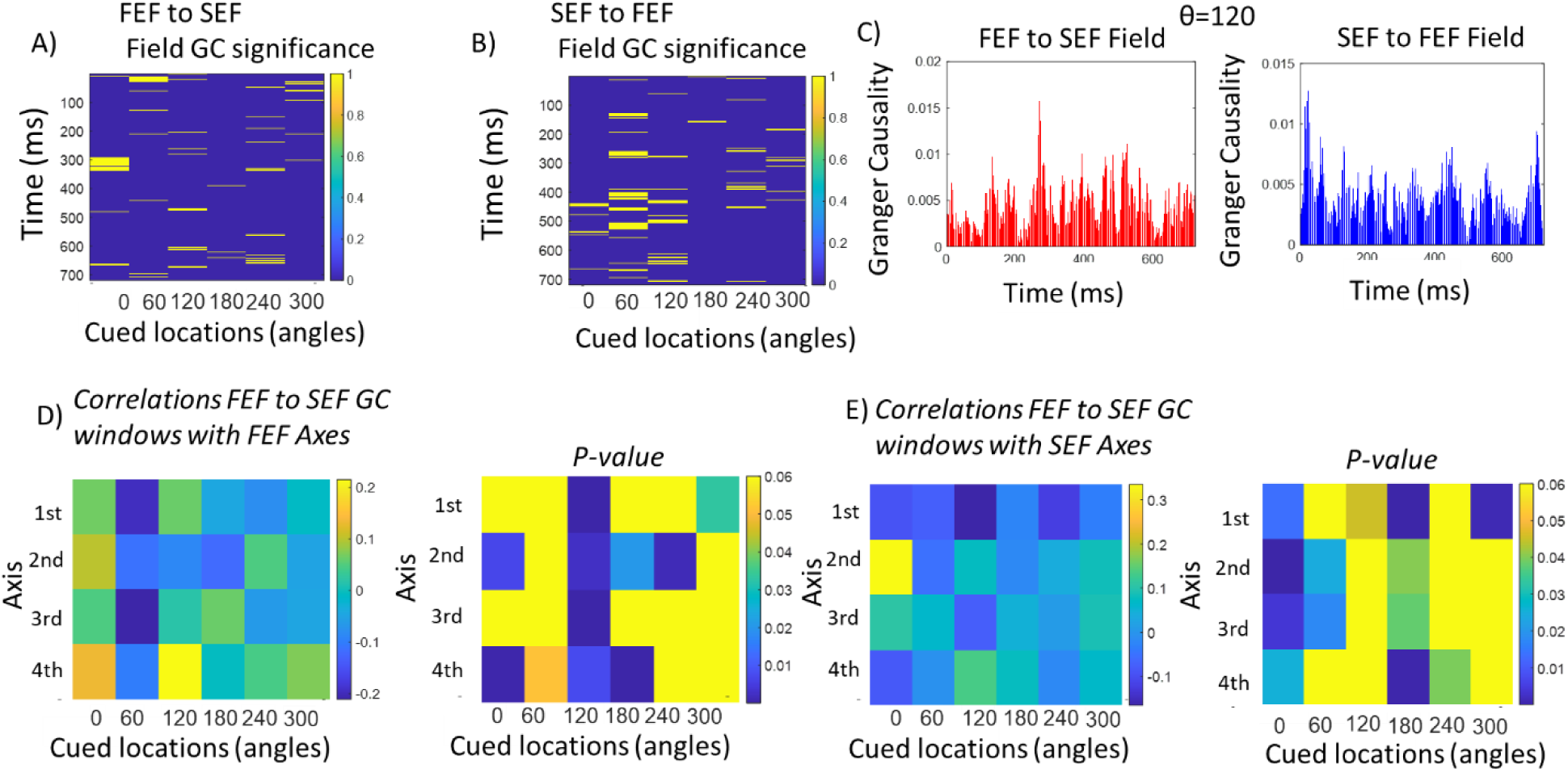
(A) Time points of significant GC field interactions from FEF to SEF for all cued locations. Time is shown on the vertical axis and cued locations on the horizontal. Significant interactions are shown in yellow. (B)Similar to 4A. Significant GC field interactions for the reverse direction, from SEF to FEF. (C)GC strengths (vertical axis) of FEF to SEF (left panel) and SEF to FEF (right panel) field interactions across time (horizontal axis) for θ = 120 degrees. (D)Correlations (left panel) and p values (right panel) between FEF principal axes and temporal windows during which GC field interactions from FEF to SEF were significant. Principal axes are shown on the vertical axis (from first to fourth as we move downwards) and cued locations on the horizontal axis.(E) Similar to Figure 4D for SEF principal axes.

**Figure 4.**
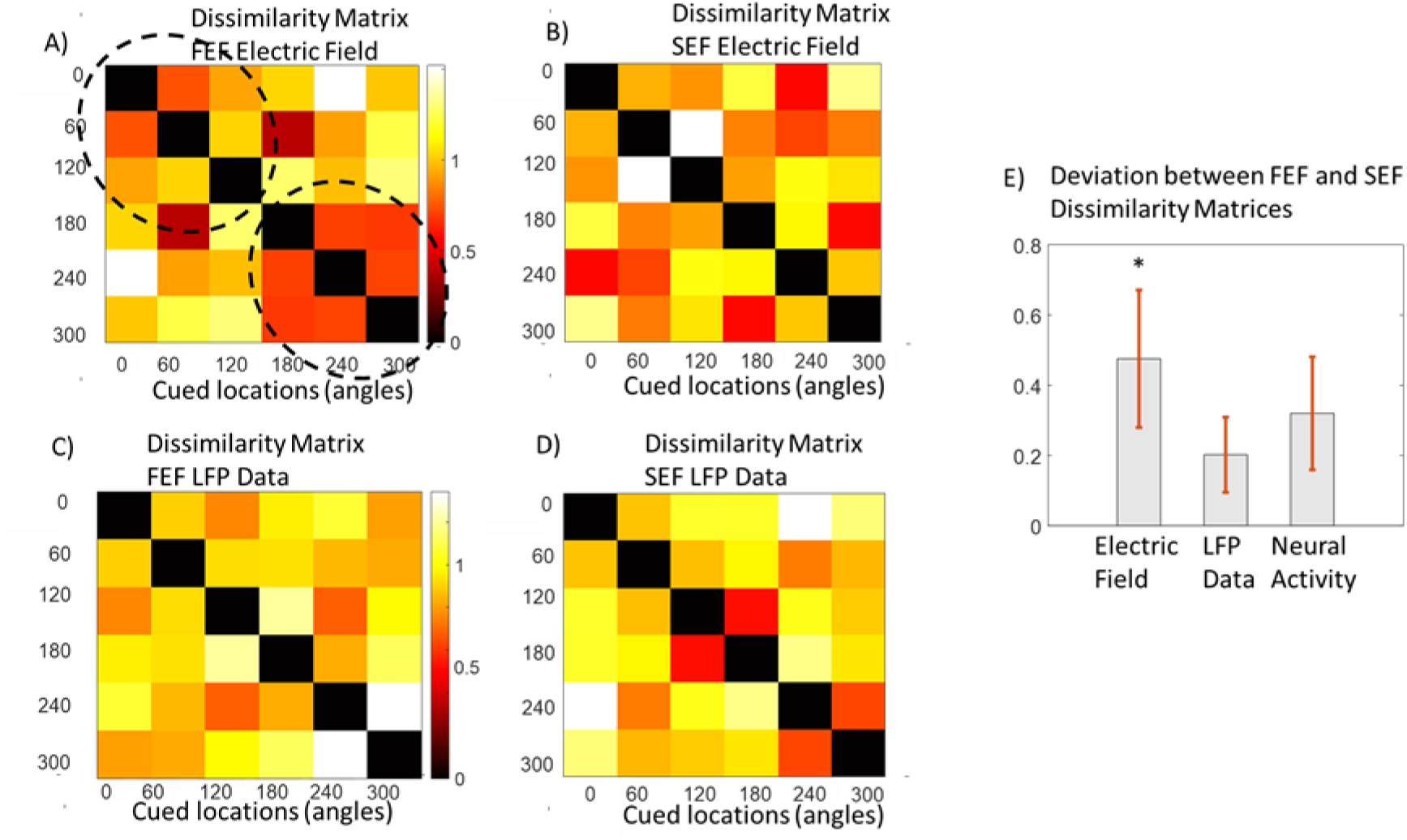
(A) Representation Dissimilarity Matrix (RDM) computed using FEF electric fields. Notice the lattice structure shown inside the dashed ellipses reminiscent of topographic clustering in FEF. (B) RDM computed using SEF electric fields. (C) RDM computed using FEF data. (D) RDM computed using SEF data. (E) Deviations (second order correlations) between RDMs. Deviation for electric field RDMs was the only that was significant (denoted by an asterisk above the leftmost bar; significance at the p<0.05 level). Error bars denote the standard errors (N=100).

Example field-to-field GC strengths for a cued location at *θ* = 120 degrees are shown in Figure 4C. FEF to SEF field GC strengths are shown in the left panel (red). GC strength in the reverse direction is shown in the right panel (blue). GC strengths are on the vertical axis. Time points are on the horizontal axis. Strengths have similar ranges in both directions during the delay period. We found similar results using neural activity (Supplementary Figure 2).

Like the results based on electric fields discussed above, Supplementary Figures 2A and 2B reveal temporal windows of information transfer between FEF and SEF at the neural activity level. Supplementary Figure 2C shows GC strengths in both directions. Interactions at the level of neural activity are expected: we found above that electric fields guide neural activity and that there were significant interactions between FEF and SEF electric fields. GC interactions at the level of neural activity are sparser than the corresponding GC strengths at the electric field level and this is replicated across all cued angles (results not shown). There are fewer red and blue lines in the left and right panels of Supplementary Figure 2C compared to Figure 4C. At several time points, GC strengths based on neural activity were zero, while GC strengths based on fields were not. This confirms the stability and robustness of the electric field found above and in our earlier work.

Are the temporal windows during which significant electric field interactions occur related to neural activity fluctuations? If so, this would mean that the dynamics (fluctuations) of neural ensembles in FEF and SEF are linked to the information transfer between them. This is what we tested next. We asked whether the temporal profile of significant field interactions found using GC above (yellow lines in Figures 4A and 4B) follows the neural dynamics in each brain area. Our hypothesis was that significant field interactions would occur while neural activity fluctuations were relatively large. We thus looked for correlations between the temporal windows (epochs) during which GC significant field interactions took place and neural activity.

Our hypothesis was that ephaptic interactions would be sensitive to both the amplitude and the spatial extent (scale) of neural activity. This is motivated by the fact that larger amplitudes would increase SNR and functional connectivity is known to be expressed within certain frequency bands. This is known as Communication-through-Coherence (CTC) hypothesis (Fries, 2015). In (Pinotsis et al., 2017b), we showed that functional connectivity in certain bands can be described by the *principal axes* of our model, see *Methods* and (Pinotsis et al., 2017b).The axes provide approximations of the fluctuations of neural activity around baseline at different spatial scales. They are matrices of dimensionality number of time samples *by* number of trials. They describe the instantaneous contribution to the recorded LFP data averaged over electrodes. To test whether there was any relation between the timings at which GC interactions occurred and neural activity, we computed the correlations between the first, second, third and fourth principal axes and the epochs during which field GC was significant. As the order increased, the spatial scale of neural activity described by principal axes, increased too.

In Figure 4D, we show correlations (left panel) and the corresponding *p* values (right panel) between FEF principal axes and temporal windows during which electric field GC interactions from FEF to SEF were significant. Principal axes are shown on the vertical axis (from first to fourth as we move downwards) and cued locations on the horizontal axis. Figure 4E includes the corresponding results for SEF principal axes. Different colors in the *p* values (right) panel correspond to different significance levels –where we have lumped together all *p* values above the significance threshold (*p=*0.05) and shown them in yellow. The same visualization is followed in Figure 4E and Supplementary Figures 2D and 2E. In brief, yellow entries denote non-significant correlations in these Figures.

Overall, for both FEF and SEF and all cued angles, the temporal windows during which FEF to SEF CG strengths were significant, correlated with principal axes, i.e. endogenous fluctuations around baseline. P*-*values in each column (cued location) in the right panels in Figures 4D and 4E includes non-yellow, i.e. significant correlations. Interestingly, this was not the case for GC strengths based on neural activity (Supplementary Figures 2D and 2E). For certain angles, there were no significant correlations between GC strengths based on neural activity and fluctuations (principal axes). This was the case for correlations with FEF axes for *θ* = 0 degrees (right panel in Supplementary Figure 2D) and with SEF axes for *θ* = 180 and 240 degrees (right panel in Supplementary Figures 2E).

Thus, we found that fluctuations around baseline activity in both areas correlated with the temporal windows of significant field GC interactions. The evolution of information transfer between FEF and SEF follows the dynamics of the neural ensembles in these areas. The link between information transfer (significant GC windows) and neural dynamics appeared stronger at the level of electric fields. The above results suggest that electric fields are more stable than neural activity.

Above we found significant interactions at the level of electric fields in both directions between FEF and SEF (Figures 4A and 4B). We also found that interactions in the FEF to SEF direction followed the dynamics of neural ensembles (Figures 4D and 4E). Interestingly, interactions for several cued locations GC strengths in the reverse direction were non-significant. Supplementary Figure 3B (left panel) shows this was the case for FEF fluctuations and cued locations at *θ* =180, 240 and 300 degrees. The right panel in the same figure shows absence of significant correlations with SEF fluctuations (SEF axes) for *θ* =120,180 and 240 degrees. This suggests that information flow in the memory network seems to follow FEF, not SEF neural ensemble activity. SEF activity at the same time, includes both information flowing out from SEF and reverberating delay activity in the FEF—SEF network.

### The same memory is stored by electric fields in different brain areas

In the previous section, we used Granger causality and found that information was transferred between brain areas, FEF and SEF, during memory maintenance. Our hypothesis was that data were recorded from sites that are part of engram complexes.

To confirm this, we asked whether representations (engrams) in each site corresponded to the same memory. To test for similarity between information content we used *Representation Similarity Analysis* (*RSA*; Kriegeskorte et al., 2008, *Methods*). First, one constructs Dissimilarity Matrices (DMs) based on correlation distance to evaluate the similarity between memory representations. DMs describe pairwise differences in patterns of neural activity or electric fields corresponding to different cued locations. In turn, correlation distances between DMs, known as deviations, express second order differences, that is, differences in pairwise differences in neural activity or electric fields in different brain areas for the same cued locations. We used deviations to test for significant correspondence between memory representations (Diedrichsen and Kriegeskorte, 2017; Peterson et al., 2018; Pinotsis et al., 2019).

We first constructed DMs for FEF and SEF based on three different sets of data: electric fields, LFPs and neural activity. Fields and activity were reconstructed using our model *(Methods)*. Our results are shown in Figure 4 and Supplementary Figure 4. Figures 4A and 4B include the DMs for FEF and SEF electric fields respectively. Figures 4C and 4D include the corresponding RDMs based on LFPs and Supplementary Figures 4A and 4B include RDMs based on neural activity. Different colors correspond to different dissimilarities (1-correlation) for each of the six possible cued locations.

Correlations were computed between trials corresponding to the same stimulus for all possible stimulus pairs after averaging over time. The higher the dissimilarity the more variability in the way information is represented. In other words, DMs illustrate the geometry of stimulus space, that is, how different cued locations are distributed into the space spanned by the activity of the underlying neural ensemble or its electric field. This provides a visualization of how dynamics in different brain areas represent memories. It can reveal clusters implying categorical representations or smooth variations along stimulus dimensions that link to behavior. The overall structure of matrices in Figures 4A and 4B describes how the electric field representations differ between pairs of cued angles. Diagonal terms have zero dissimilarity as expected. Representations were different between stimuli (red and yellow entries, *d* ≥.4). This is also the case for other RDMs in Figures 4C and 4D as well as Supplementary Figures 4A and 4B. Interestingly, FEF RDMs based on electric field and neural activity Figure 4A and Supplementary Figure 4A show a lattice structure: representations corresponding to the upper (*θ* = 0,60 and 120 degrees) and lower (*θ* =180, 240 and 300 degrees) hemifield form distinct clusters, shown by ellipses. This is reminiscent of topographic clustering in FEF, that is known to contain topographically organized responses and visual map(Funahashi et al., 1989; Thompson and Bichot, 2005). It is also in accord with a similar organization of functional and effective connectivity found using the same dataset in (Pinotsis et al., 2017a).

To confirm that representations contained the same memories, we then computed the deviations between DMs (Kriegeskorte et al., 2008; Pinotsis et al., 2019). Our results are shown in Figure 4E. Deviation is a second order correlation distance, that is, the distance between correlation distances shown in DMs. It allows us to quantify matches between memory representations in the two areas. The smaller the deviation the closer the match. To test whether two DMs were related, we used fixed effects randomization test. We simulated the null distribution by reordering rows (10,000 relabelings) and obtained a distribution of correlations (the null hypothesis is that the two DMs were unrelated). If the actual correlation we had obtained fall within the top 5% of the simulated null distribution, then we reject the null hypothesis: the two DMs are related. Figure 4E shows that the deviation for electric fields was larger (*d* ≍ .5) than that computed using neural activity (*d* ≍ .3) which, in turn, is larger than the deviation computed using LFPs (*d* ≍ .2). Crucially, the randomization test reveals that only the DMs based on electric fields are significantly related (denoted by an asterisk above the leftmost bar in Figure 4E; significant deviations at the *p<*0.05 level). The significant relationship between DMs based on electric fields suggests that memory representations contain unique information associated with different memories. This is in accord with earlier results from (Pinotsis and Miller, 2022) obtained using the same data, which found that classification accuracy was higher when electric fields were used as features compared to neural activity. They also found that confusion matrices based on fields had more correctly classified trials. In Figure 4E, error bars denote the standard errors. They depict the variability of deviations (had we chosen different stimuli from the same population; *N=*100, see (Kriegeskorte et al., 2008)).

To sum, we found significantly related dissimilarity matrices in FEF and SEF computed using electric fields, but not LFPs or reconstructed neural activity. This suggests that memory representations in the two areas, known as engrams, are linked at the electric field level. Crucially, these similarities in memory representations across two areas were not apparent in LFP recordings. Taken together with our earlier result that emerging electric fields seem to guide information transfer, our result here suggests that electric fields mediate the transfer of memories and their latent states between brain areas. Ephaptic interactions occur in areas where engrams are found. See *Methods* for mathematical arguments supporting this result.

## Discussion

We found evidence for in vivo ephaptic coupling from two cortical areas, the FEF and SEF, during performance of a spatial delayed response task. We found that ephaptic coupling from bioelectrical fields is causative, it influences neural activity, sculpting and guiding it to form engram complexes. We found that, in each brain area, information was transferred from bioelectric fields to neurons. Also, stable, robust fields allowed for memory transfer between FEF and SEF engrams. Neural activity appeared to contain less information and was more variable. In short, like a conductor of an orchestra, where neurons are the musicians, the bioelectric field influences each neuron and orchestrates the engram, the symphony.

To demonstrate ephaptic effects, we used biophysical modelling and Granger causality. We used a model that can describe neural ensemble connectivity, synaptic filtering, and electric fields. In previous work, we estimated the effective connectivity in neural ensembles and their electric fields (Pinotsis et al., 2017a; Pinotsis and Miller, 2022a). We found that electric fields carry stimulus information, are robust and can act as “guardrails” that stabilize and funnel the underlying neural activity. We showed that fields were more stable than neural activity and could be used to decode remembered cued locations better.

Here, we used the same model and tested whether including ephaptic effects resulted in better fits to LFP data. The model was used for both learning and inference. It first learned the connectivity parameters. These were subsequently used as priors to reconstruct single trial neural activity and bioelectric field estimates. This revealed ephaptic effects when endogenous fluctuations were small, as expected from the linearity assumption of our model. Granger Causality (GC) applied to time snapshots confirmed these ephaptic effects during large endogenous fluctuations and also allowed us to determine directionality.

Our results were consistent with the communication through coherence hypothesis (CTC (Fries, 2015)). According to CTC, neural ensembles synchronize in a way that creates bursts of excitation and inhibition and allows information to propagate from one area to the other during certain temporal windows. We took CTC one step further to suggest that this communication is guided by emerging electric fields. First, we found that between area GC strengths based on fields were larger than the corresponding estimates based on neural activity. Second, for each brain areal GC strengths were much larger in the field-to-activity than in the reverse direction. Third, the temporal windows during which FEF to SEF interactions take place followed the dynamics of neural ensembles in these areas. Taken together, the above results suggest that electric fields guide information transfer between areas.

The last result, that fluctuations around baseline in FEF and SEF correlated with the temporal windows of significant field GC interactions suggests a circular causality between neural sources and emerging fields. Had we measured interactions with the ordinary GC from time series analysis, one would expect GC to be stronger when neural activity increased because of increased SNR. However, we here considered instantaneous interactions. Electric field effects on neurons travel at the speed of light. Thus, interactions do not depend on synaptic and conduction delays that would be measures with ordinary GC. We used a different measure, “spatial GC” to describe interactions and calculated GC strengths based on time snapshots or, in other words, spatial profiles of neural activity —instead of time series. Finding that large fluctuations in neural activity correlated with windows of significant spatial GC interactions suggests that neurons generated electric fields that fed back to them instantaneously. This is a form of circular causality. Circular causality is central in the theory of synergetics discussed below.

We also found that the electric fields were more stable than neural activity, i.e., had less representational drift. The coefficients of variation associated with field -to-activity GC strengths were smaller than the corresponding coefficients based on activity-to-field interactions. Also, GC strengths of interactions between FEF and SEF neural activity were sparser over time than the corresponding strengths based on electric fields. This concurs with earlier results where electric field estimates were more often correlated across trials, i.e. more stable, compared to neural activity estimates (Pinotsis and Miller, 2022a).

Using Representation Similarity Analysis (RSA; Diedrichsen and Kriegeskorte, 2017; Kriegeskorte et al., 2008)), we also confirmed that electric fields emerging from FEF and SEF ensembles contained the same information. RSA assesses matches between memory representations in different brain areas. Information can be represented at different levels, e.g. in neural activity or electric fields. We found that FEF and SEF representations contained similar information only when we used electric field data for RSA analysis – not LFP or neural activity. Thus, memory representations seem to be linked at the electric field level.

Overall, our results suggest that in addition to synaptic transmission, information transfer might be guided in a top-down fashion by electric fields. In mathematical language, electric fields are a control parameter. This term appears in the theory of synergetics from complex systems (Basar et al., 2012; Haken, 1987). Examples of control parameters include energy (Basar et al., 2012; Haken, 2012), and feedback attention signals in a binocular rivalry task (Ditzinger and Haken, 1989). A control parameter has two features that the electric field has: it is stable and evolves at a slower time scale than enslaved parts (i.e. neural activity). In *Methods*, using mathematical arguments, we explained these features and showed how ephaptic coupling follows from the slaving principle (Haken, 2012 ; see also discussion below). We also showed that if ephaptic coupling occurs in one brain area in a memory network (engram complex) it will occur in all other brain areas.

Our results offer a plausible explanation of ephaptic coupling as an application of the more general slaving principle of synergetics. Of course, other explanations of the slow dynamics of emerging electric fields might exist. For example, synaptic plasticity or slow waves of synaptic barrages could also play a role. We will consider this in future work.

The idea that electrical fields play a role in the formation of neural ensembles has a long history. The connection between memories, connectivity and electric fields was noted early. The term engram complex was coined by German biologist Richard Semon, who, over a century ago, suggested that memories are stored in groups of neurons in multiple brain areas (Semon et al., 2018). Then, according to Semon’s law of ecphory, memory recall happens when an appropriate electric field is generated —an energetic “condition” similar to memory registration is achieved during recall (Josselyn et al., 2015; Semon et al., 2018; Thompson, 1976).

The importance of the electric field has also been emphasized in recent synaptic plasticity studies. These have revealed that learning and memory change scaffolding proteins that regulate synaptic functions, like trafficking and binding of NMDA or other receptors (Kim and Sheng, 2004). In turn, protein changes result in changes of synaptic activity and of the electric field in the extracellular space. Thus, synaptic activity is not dictated solely by electrical elements, the receptors, charged particles and currents, but also chemical elements, like scaffold proteins. Both electrical and chemical elements determine the electric field in the extracellular space (Queenan et al., 2017). Receptors occupy synapses with some probability, and can vary from trial to trial where the same memory is recalled. This also means that different neurons form ensembles in different trials where the same memory is maintained, a phenomenon known as representational drift(Rule et al., 2019).

It is now known that the brain’s endogenous electric field feeds back to the activity of individual ion channels and alters their neuronal firing, i.e., there is ephaptic coupling (Anastassiou et al., 2011), see (Anastassiou and Koch, 2015) for a review. The pioneering study by Eccles and Jaeger (Eccles and Jaeger, 1958) showed ephaptic effects on ion currents in synaptic cleft. McFadden and other authors have taken the importance of ephaptic coupling one step further: they have linked it to conscious awareness and hypothesized that it can be used for computation that occurs momentarily and is distributed over space (Fingelkurts et al., 2012; John, 2005; McFadden, 2020; Pockett, 2000). Direct evidence of ephaptic coupling has been found in slices (Anastassiou and Koch, 2015; Chiang et al., 2019; Jefferys et al., 2012). Testing such hypotheses and in vivo ephaptic effects in general is more difficult. Electrodes are far from the neural ensemble and multiple groups of neurons are activated at the same time. Further, chemical processes like electrodiffusion and others alter the electric fields (Savtchenko et al., 2017). Here, we used a variety of computational techniques to provide in vivo evidence.

The low-dimensional stability of electric fields can help the brain with memory maintenance and cognitive processing in general. Synergetics suggests that latent states, like connectivity, can be reliably transferred between brain areas, in accord with modern engram theory (Ryan et al., 2015). This is orchestrated by control parameters. In synergetics, latent states are called order parameters (Gallego et al., 2020; Yu et al., 2008). The theory posits that order and control parameters exist in all self-organized dynamical systems (e.g. molecules, fluids) and therefore the brain. They emerge because of self-organization and capture collective dynamics of a large part of the system’s individual parts. Importantly, parts, order and control parameters evolve at different timescales that are separate: control (bioelectric fields, slowest), order parameters (e.g. effective connectivity, oscillation frequency, intermediate) and enslaved parts (spiking, fastest).

This separation of timescales follows from the center manifold theorem. Haken (Haken, 2006) pointed out this separation is crucial for consciousness. Order parameters evolve slowly and this “can be interpreted as a phase transition from subliminal to conscious phase”. They sent essential information to other brain areas. This is like Mooney faces that are images known to induce gamma oscillations associated with conscious experience (Lachaux et al., 2005). Oscillation frequency is an order parameter in this case. In (Pinotsis et al., 2018), we showed that during a working memory task, when the cognitive capacity limit was exceeded, synchrony between oscillatory responses in PFC, FEF and LIP broke down and the monkey made errors. That is, order parameters were different when the monkey could vs. when he could not remember. More generally, neural ensembles are thought to maintain memories as a result of coordinated neuronal activity (Tonegawa et al., 2015b). Control parameters can control the spiking of such large numbers of neurons. We here suggest that the electric field is a control parameter. Control parameters guide order parameters and constrain enslaved parts. Neurons give rise to the ensemble and an emerging electric field. This, in turn, determines the function of each neuron through ephaptic coupling. This is an example of circular causality and an application of the slaving principle mentioned above.

This is also a difference between synergetics and dimensionality reduction approaches (Gao and Ganguli, 2015; Jazayeri and Ostojic, 2021). Like dimensionality reduction, synergetics uses latent states. But it also uses control parameters. These evolve at an even slower timescale than latent states and spiking and are characteristic for each state of the brain, e.g. each memory. Synergetics suggests that control parameters are somehow fixed in the sense that when they change, the brain goes to a different stable state, similar to phase transitions in thermodynamics (Domb, 2000).

In sum, using biophysical modeling, machine learning and Granger causality we provided some evidence supporting the hypothesis that bioelectric fields are a control variable that enslaves neural activity. This can have implications for modern BCI, where electric field manipulations are used to control neurons so that activity reverts to a healthy state and patient behavior is abolished.

## Acknowledgements

This work is supported by UKRI ES/T01279X/1, Office of Naval Research N00014-22-1-2453, The JPB Foundation and The Picower Institute for Learning and Memory.

## Footnotes

1 LFP electrodes measure potentials *V^e^*. Thus, we can assume that the location of the LFP electrode denoted by a star in Figure 1B coincides with the point *P*(*x*, *y*) where the electric field is evaluated.

2 As expected the complexity of both models was very similar, see Supplementary Figure 2C which shows the difference in complexity between models. All estimates are between 0-0.6, which is less than 0.5% of the BF factor shown in Figure 2A. Note also that for *θ* = 0 degrees the non ephaptic model was more likely.

3 Briefly, it explained 40% of the variance and found the same connectivity for ensembles maintained cued locations, see above.

